# Microplastics from disposable paper cups are enriched in the placenta and fetus, leading to metabolic and reproductive toxicity during pregnancy

**DOI:** 10.1101/2024.01.20.576431

**Authors:** Qiong Chen, Chen Peng, Haoteng Xu, Zhuojie Su, Gulimire Yilihan, Xin Wei, Yueran Shen, Chao Jiang

**Affiliations:** Zhejiang Provincial Key Laboratory of Cancer Molecular Cell Biology, Life Sciences Institute, Zhejiang University, Hangzhou, Zhejiang 310058, China; State Key Laboratory for Diagnosis and Treatment of Infectious Diseases, National Clinical Research Center for Infectious Diseases, First Affiliated Hospital, Zhejiang University School of Medicine, Hangzhou, Zhejiang 310009, China; Center for Life Sciences, Shaoxing Institute, Zhejiang University, Shaoxing, Zhejiang 321000, China; School of Life Sciences, Westlake University, Hangzhou, Zhejiang 310030, China

## Abstract

The health implications of microplastics (MPs), especially those originating from hot drinks in disposable paper cups (DPCs), are increasingly alarming. We investigated the accumulation and metabolic and reproductive toxicological effects of MPs from DPCs filled with hot water in various tissues in a pregnant mouse model. Simulating human intake of 0.3, 3.3, and 33.3 cups daily, we found MPs exposure-induced dose-responsive harmful effects on murine fetal development and maternal physiology. MPs were detected in all 13 examined tissues, with the highest accumulation in the cecal contents, followed by significant depositions in the fetus, placenta, kidney, spleen, lung, and heart. A higher proportion of smaller MPs (90.35% < 10 μm) was identified in brain tissues. Dose-responsive changes in functional microbiome and gene pathways were observed. Moderate MPs intake of 3.3 cups daily significantly altered cecal microbiome composition and metabolic functions. The transcriptomic functional variations in maternal blood, placenta, and mammary gland underscore the significant impacts of realistic MPs exposure on metabolic and immune health and posing neurodegenerative and miscarriage risks. The benchmark dose framework analysis using tissue-specific gene biomarkers revealed safe exposure limits at 2 to 4 cups/day during pregnancy. Our results indicate selective tissue accumulation and potential metabolic and reproductive toxicities of MPs at exposure levels presumed non-hazardous. Such risks remain unaddressed within current food safety regulations, impacting vulnerable groups such as pregnant women and fetuses.

**Research Highlights:**

- Microplastics released from disposable paper cups filled with hot water showed preferential accumulations in the murine fetus, placenta, kidney, spleen, lung, and heart, with significant adverse impacts on fetal development.
- Microplastic exposure led to dose-responsive maternal microbiome changes associated with increased fatty acid biosynthesis and elevated expressions of genes related to viral infections, neurodegenerative diseases, oxidative stress, and miscarriage risk.
- A consumption level of 3.3 cups/day was sufficient to elicit systemic metabolic and reproductive toxicity, with a predicted safe exposure limit of 2 to 4 cups/day during pregnancy by benchmark dose framework analysis with molecular biomarkers.

## Introduction

Plastic pollution significantly impacts environmental and human health. The extensive production and consumption of plastics, along with inadequate waste management, have resulted in billions of tons of plastic waste in ecosystems (*1*). Notably, 79% of plastic products are improperly treated, leading to plastic waste accumulations in landfills or natural environments (*2*). These plastics leach toxic chemicals (*3*), harm wildlife, and pose health risks to humans (*4*). Furthermore, plastics can act as vectors and reservoirs for persistent organic pollutants, heavy metals, and antibiotic-resistant microorganisms (*5–7*).

In addition, plastics undergo environmental degradation through chemical, physical, and biological reactions, with UV radiation primarily initiating this process. The degraded forms of plastic fragments, when smaller than 5 mm, are termed microplastics (MPs) (*8*). Primary MPs, intentionally added to consumer products, such as cosmetics and detergents, paints, medications, nappies, and insecticides, contribute further to this issue (*9*). Human exposure to MPs occurs via ingestion of contaminated food and water, use of personal care products (toothpaste, face wash, scrubs, soap; also, dermal route), and inhalation, leading to bioaccumulation in various tissues (*10*), including the lungs (*11*), bloodstream (*12*), and placenta (*13*). Traditional detection methods, including stereomicroscopy, SEM, FTIR, Raman, and fluorescent tagging with Nile Red, often fail to fully capture the presence and particle size distribution of MPs in biological samples (*10*, *11*, *13*, *14*).

The toxicological effects of MPs in cell cultures, organoids, and animal models include oxidative stress, DNA damage, altered metabolism, and neurotoxicity (*15–17*). The reproductive and developmental impacts of MPs are alarming, with potential intergenerational effects. For instance, Bisphenol-A (BPA) has been linked to impaired sperm maturation (*18*), and polystyrene exposure to reduced ovarian health (*19*). Parental exposure to polyethylene MPs in mice has shown reproductive and developmental issues in offspring (*20*), with rodent studies indicating liver damage and metabolic disruptions over generations (*21*). There is also evidence of intrauterine growth restriction linked to MP exposure (*22*). Despite these findings, present studies often utilize single-size and single-property MPs, instead of realistic and complex mixtures of MPs encountered in real-life (*19–21*). Many studies have administered MPs by incorporating them into drinking water, leading to imprecise MPs intake (*21*, *23*). Importantly, the concentrations of MPs used in previous studies were often much higher than regular human consumption levels. Therefore, it was impossible to extrapolate a human-health-based threshold value for MPs (*24*). Additionally, the distribution and size variation of mixture MPs deposited in different biological tissues of an organism have not been systematically investigated.

A significant source of MPs is disposable paper cups (DPCs) used for hot beverages. The release of MPs varies with beverage type, pH, and temperature, with notable differences between carbonated beverages and soda water (*25*, *26*). In 2018, China and the United States consumed approximately 95,890 and 19,178 paper cups every minute, totaling 50 and 10 billion cups per year (*27*). A recent survey indicated that 74% of Americans drink coffee daily, with 49% of people consuming 3 to 5 cups per day (https://www.driveresearch.com/market-research-company-blog/coffee-survey), raising concerns about the intake of MPs released from DPCs (*14*).

Our study began by examining DPCs from five popular anonymous brands to identify the brand with the highest MPs release. We collected and enriched these MPs, and administered them to pregnant mice through oral gavage, simulating human daily consumption levels. We evaluated MPs bioaccumulation in 13 types of tissues and found significant preferential bioaccumulation of MPs in the fetus, placenta, kidney, spleen, lung, and heart. The MPs exposure resulted in fetal growth restriction and compromised development of mammary glands in the dams. We further analyzed the microbiome of the cecal samples and transcriptomes of blood, placenta, and mammary glands from dams. Our results indicate significant dysbiosis in the cecal microbiota, even at the moderate consumption of MPs. Systemic disruptions in gene expressions related to metabolism, neurodegenerative diseases, immune responses, and miscarriage were identified. Our study sheds light on the physiological and molecular health outcomes after exposure to MPs from DPCs at realistic level, underscoring the need for further evaluating and addressing these potential health hazards with relevant food safety regulations, especially for populations at risks.

## Results

### Size distribution and characteristics of concentrated MPs from disposable paper cups

We acquired 500 ml DPCs from 5 brands and filled them with 85–90 °C ultrapure water for 15 mins (*14*) to simulate the release of MPs during consumption of hot drinks. Released MPs were observed in all brands by fluorescence microscopy images (Supplementary Fig. 1a). No MPs were detected when room-temperature ultrapure water was added, confirming that hot water is required for MPs to release (*14*). We then assessed the number of MPs released and found Brand C released the most MPs— approximately 8.02 ± 1.87 million particles/100 ml of water (Supplementary Fig. 1).

We next used 500 ml DPCs from Brand C to prepare MPs samples equivalent to human consumption of 0, 1, 10, and 100 cups every three days and labeled the groups as 0C, 1C, 10C, and 100C accordingly (Fig. 1a, b). It is important to note that 1 and 10 cups of hot beverages every three days are very reasonable human consumption levels in modern societies (*28*). Fluorescence analyses by BioTek Cytation 3 (Methods) evaluated the number and size distribution of MPs in each group (Fig.1c). The particle sizes of 1C, 10C, and 100C groups ranged from 6 - 70 μm, 6 – 62.1 μm, and 6 – 154 μm, with 60.33%, 58.21%, and 59.27% of the particles under 10 μm, respectively (Fig. 1d, e). All treatment groups showed similar median particle sizes (Fig. 1e). For precise quantification, MPs of each sample were analyzed using the BioTek Cytation 3 plate reader in a 96-well plate. Cytation 3 is a cell imaging multi-mode microplate reader that combines automated digital microscopy and conventional microplate detection. This instrument can cover entire well with 28 images in less than 5 minutes. To ensure accuracy, each 96-well plate included three processing blank controls to correct for background fluorescence. This methodology accurately detects MPs as small as 6 μm in diameter. We applied the same acquisition and analysis parameters across all wells and plates.

**Fig. 1.**
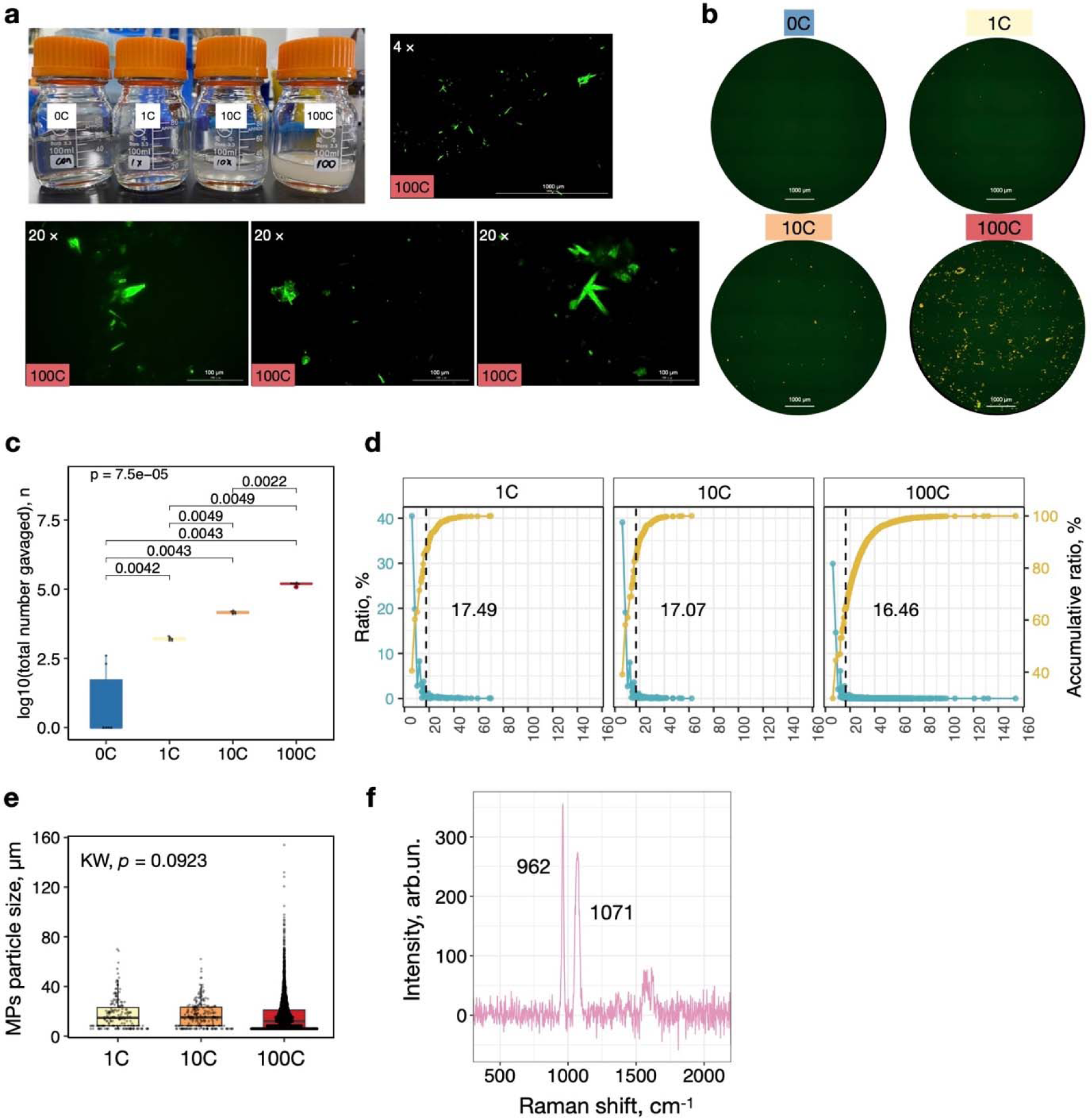
Size distribution and characteristics of concentrated MPs from DPCs. (a) Enriched MPs prepared for four exposure groups, with representative images of the 100C group at 4x and 20x magnifications. (b) Representative fluorescent composite images of enriched MPs from all groups, each composed of 28 images using the montage function and captured at 4x magnification. (c) The total number of MPs administered per dam during gestation, with statistical differences assessed using the Kruskal-Wallis test with post hoc Games-Howell tests. (d) Particle size frequency distribution (blue) and cumulative distribution (yellow) for the 1C, 10C, and 100C groups. Vertical blank dashed line indicates the average particle sizes. (e) Boxplots illustrating the size distribution of MPs in the 1C, 10C, and 100C groups, p-values calculated by Kruskal-Wallis tests. (f) Raman spectroscopy profiles of MPs extracted from DPCs.

To evaluate the MPs’ properties, we performed Raman Microspectroscopy on the 100C group in triplicate. The resulting spectra were matched against the KnowItAll, SLOPP, and SLOPPE databases (Methods). The Raman signatures were consistent with polystyrene (Fig. 1f), highlighted by the dominant peak at approximately 1000 cm^-1^ due to the aromatic ring structure, indicative of π-π stacking interactions and the characteristic ring-breathing vibration band (*29*).

### MPs exposure led to maternal physiological changes and fetal growth restrictions

To study the toxicity of DPC-derived MPs, we administered different dosages of MPs in pregnant murine models through gavage and subsequently analyzed the developmental and physiological outcomes (Fig. 2a). Post-exposure assessments revealed no significant alterations in the maternal body weight (Supplementary Fig. 2a). Mammary gland development was quantitatively evaluated through alveolar count, alveolar area, and the ratio of alveolar area to gland area. MPs exposure had no significant impact on alveolar count or area (Supplementary Fig. 2b, c); however, a negative correlation with the alveolar ratio was significant in the generalized linear model (GLM), revealing a dose-dependent relationship (Fig. 2b). The GLM approach extends the capabilities of linear regression to accommodate response variables with non-normal error distributions. In our model, we treated the MP dose as continuous independent variables with values of 0, 1, 10, and 100 to model the regression of doses against observed outcomes. Additionally, an increasing trend in relative liver weight in the MPs-exposed dams indicate potential liver hypertrophy or hyperplasia (Fig. 2c).

**Fig. 2.**
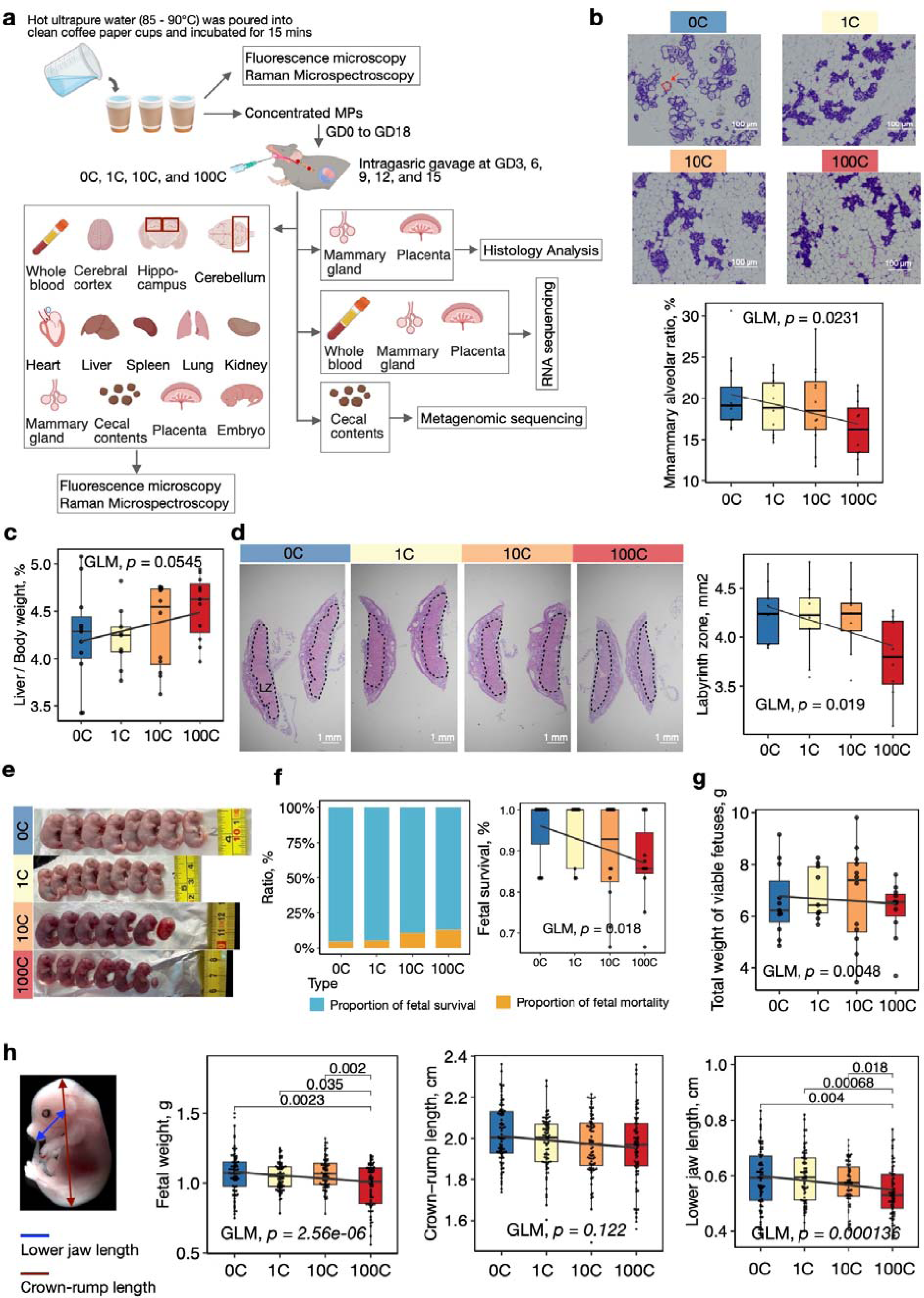
The experimental design and impact of maternal MPs exposure on maternal and fetal physiology. (a) The overall experimental design. (b) Hematoxylin and eosin (HE) staining (top) and quantification of mammary alveolar percentage (bottom), with p-values of GLM. The N in each group were 0C (13), 1C (11), 10C (12), and 100C (11). (c) Ratio of liver to body weight, with p-values of GLM. (d) HE staining (left) and quantification of placenta labyrinth zone (right), with p-values of GLM. (e) Representative images of litter in different treatment groups. (f) Rates of fetal survival, with p-values of GLM. (g) Total weight of fetuses in each litter, with p-values of GLM. (h) Assessment of fetal growth impact from maternal MPs exposure. The above p-values with statistical differences were assessed using the Kruskal-Wallis test with post hoc Games-Howell tests, besides p-values from GLM.

Furthermore, we incorporated litter size as a covariate in the GLM to examine the relationship between MP doses and fetal growth parameters. A dose-dependent reduction in the placental labyrinth zone was noted, indicative of compromised placental function (Fig. 2d). This impairment correlated with a reduction in fetal viability in a dose-dependent manner (Fig. 2e, f). The total weight of viable fetuses (Fig. 2g) and individual fetal metrics, such as weight and lower jaw length, also showed a negative dose-dependent correlation (Fig. 2h).

Collectively, these findings underscore the potential risks associated with maternal exposure to MPs from DPCs, manifesting as altered mammary gland development, dose-dependent constraints on fetal growth and development, and potential changes in liver physiology.

### Heterogeneous distribution and size variation of MPs in fetal-maternal tissues

Systematic investigation of the deposition of MPs across an array of tissues within a single organism has not been documented before. Utilizing the BioTek Cytation 3, we quantitatively analyzed MPs size distribution in thirteen distinct biological samples of each mouse, encompassing a range of maternal and fetal tissues, as well as cecal contents, in triplicate. Notably, the absence of MPs in blank controls indicate that our approach is robust (Supplementary Figs 3, 4). A clear dose-response relationship emerged, as most tissues exhibited an increased MP count corresponding to higher exposure levels, with the notable exceptions of the hippocampus and lungs (Fig 3a, and Supplementary Figs 3, 4).

**Fig. 3.**
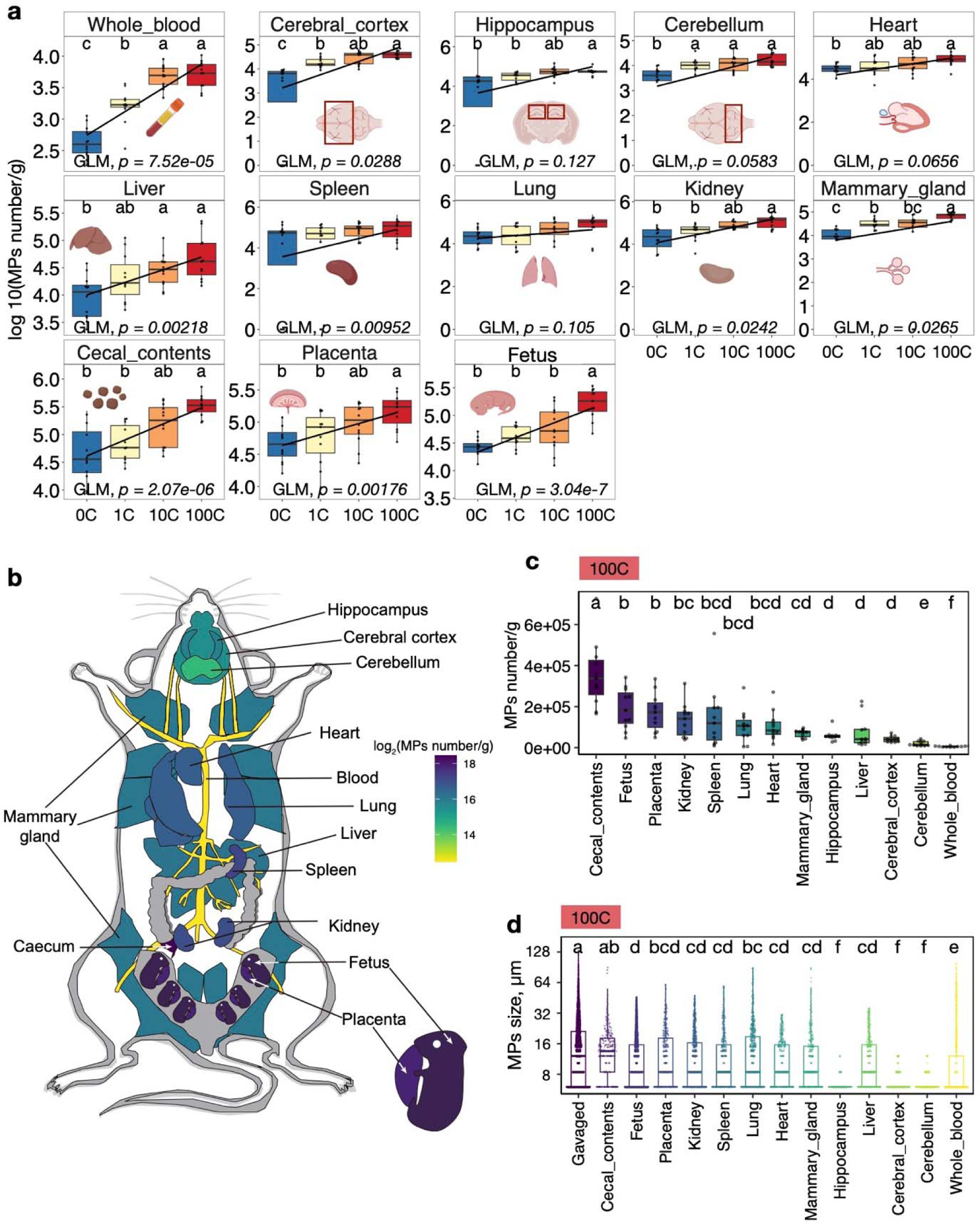
MP depositions in 13 types of biological samples. (a) Normalized number of microplastics in various biological samples, with statistical significance denoted by different lowercase letters for p < 0.05 via Kruskal-Wallis test with post hoc Games-Howell tests. GLM statistics also shown. (b) Overlay of MPs deposition on a mouse anatomical diagram; the colored gradient represents the varying density of MPs across different tissues. (c) MPs accumulated in different types of tissues in the 100C group, with significance indicated by lowercase letters for p < 0.05 via Kruskal-Wallis test with post hoc Games-Howell tests. (d) Size distribution of MPs in the 100C group, with statistical differences marked by lowercase letters for p < 0.05 as determined by the Kruskal-Wallis test with post hoc Games-Howell tests.

Among thirteen biological samples, blood samples consistently exhibited the lowest accumulation of MPs (Fig. 3b, c and Supplementary Fig. 5a, b), indicating that MPs in the blood are taken up by other types of tissues. In the 100C exposure group, the cecal contents demonstrated the most substantial concentration of MPs, reaching 41,396 ± 14,570 MPs per gram of tissue. Strikingly, we observed higher amounts of MPs in the fetus (190,436 ± 99,103 n/g), placenta (171,573 ± 92,071 n/g), kidney (134,695 ± 82,546 n/g), spleen (148,480 ± 157,691 n/g), lung (105,119 ± 80,522 n/g), and heart (105,827 ± 71,088 n/g). The mammary gland and liver also showed noteworthy MP deposition, with 69,192 ± 19,374 n/g and 72,746 ± 74,904 n/g, respectively. Notably, brain tissues were not precluded from MP deposition. This distribution pattern was consistent in the 10C exposure group as well (Fig. S5a). In the 1C group, a uniform distribution of MPs was observed across the placenta, cecal contents, spleen, kidney, fetus, heart, mammary gland, and liver, indicating a different deposition mechanism at the lower exposure level (Supplementary Fig. 5b). These findings suggest the preferred bioaccumulation of MPs in fetal and placental tissues when subjected to higher amounts of MPs exposures.

In addition to the variations in quantity, we observed significant variations in the size of MPs across different biological tissues (Fig. 3d and Supplementary Fig. 5c, d). In the 100C group, among the 13 types of biological samples, most MPs were smaller than 10 μm, accounting for 55.16% to 90.73 % of detected particles (Fig. 3d and Supplementary Fig. 6a). Interestingly, the deposited MPs were generally smaller than the administered MPs, except for those in the cecal contents, suggesting a potential sifting effect during GI absorption or breakdown processes in different tissues (Fig. 3d). The similar particle size distribution observed in the fetus and placenta indicates limited sifting by the placenta, so MPs can get directly into the fetus. Interestingly, MPs found in the 10C and 1C groups within biological samples were smaller than in the 100C group, which can be attributed to the absence of administrated larger MPs (70 to 154 μm) in these groups (Fig. 1d and Supplementary Fig. 5c, d). The smallest MPs were detected in brain tissues across all dosages, likely due to the blood-brain barrier (BBB) (Fig. 3d and Supplementary Fig. 5c, d). The Raman spectroscopic analysis confirmed that these particles are similar to the ingested MPs (Supplementary Fig. 6b).

### Microbiome functional disturbances in response to MPs exposure

To investigate the impact of MPs exposure on the gut microbiome as a possible mechanism underlying the phenotypes, metagenomic sequencing was conducted on DNA extracted from cecal contents of dams from all exposure groups. Compared to other regions of the mouse gut, the cecal microbiome is the optimal representation of the gut microbiome affected by environmental factors (*30*). Utilizing the Illumina NovaSeq 6000 platform, we generated 1.85 billion reads for 47 samples, post-removal of host DNA. We used Kraken2 (*31*) and a comprehensive reference genome database for classification, covering over 431,109 species across all life domains (Methods). We identified 145 phyla, 492 genera, and 1080 species (Fig. 4a). For subsequent compositional analysis, we used the hyperbolic arcsine-transformed counts per million (aCPM) (*32*, *33*).

**Fig. 4.**
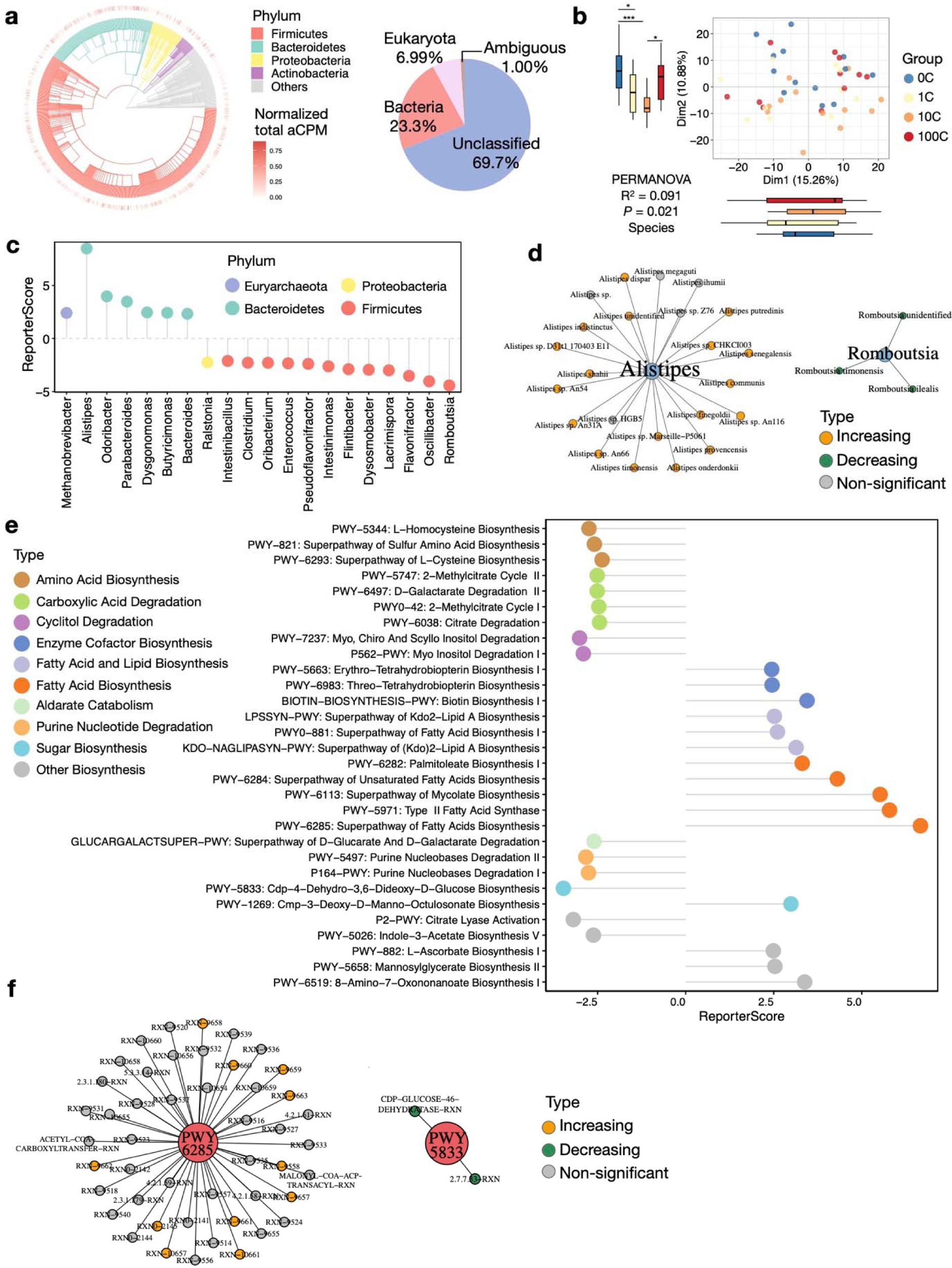
Delineating microbiome composition and functional alterations. (a) Microbiome diversity presented with an outer ring indicating the aCPM of each species. The pie chart details the total relative abundance of microbial groups. (b) PCoA analysis of samples at the species level, with boxplot showing significance denoted by asterisks as determined by Kruskal-Wallis test (***p < 0.001; **p < 0.01; *p < 0.05). (c) Lollipop chart presenting dose-dependent, significantly enriched genera (|ReporterScore| > 2.5, p < 0.05). (d) Network plot illustrating abundance changes in genera *Alistipes* (increases with MP exposure; orange) and *Romboutsia* (decreases with MP exposure; green). (e) Lollipop chart of significantly enriched pathways (|ReporterScore| > 2.5, p < 0.05) across functional categories. (f) Network plot of PWY6285 (increases with MP exposure) and PWY5833 (decreases with MP exposure).

Principal Coordinates Analysis (PCoA) based on Bray-Curtis distances between the microbiome profiles showed that different exposure groups diverged along Dim 2 (Fig. 4b). Interestingly, the 10C group (roughly 3 cups/day) appeared to be the most distinct (Fig. 4b). To elucidate dose-responsive taxonomic shifts, we performed taxonomic enrichment analysis using the genus-species relationships with the Reporter Score Analysis (RSA) method (*34*, *35*). At the genus level, the most significant dose-dependent changes were observed in the phyla of Firmicutes and Bacteroidetes (Fig. 4c), with notable increases in Bacteroidetes and decreases in Firmicutes species. However, we did not observe a significant shift in the overall Firmicutes/Bacteroidetes ratio across exposure groups (Supplementary Fig. 7a). *Alistipes* was the top increasing genus in response to MP exposures (Fig. 4d and Supplementary Fig. 7b) and are associated with inflammation and cancer (*36*). In contrast, *Romboutsia* was the top decreasing genus (Fig. 4d and Supplementary Fig. 7b), which is associated with gut homeostasis and was potentially deprived in malnutrition (*37*) and cancer (*38*). The variations in the cecal microbiome suggested impaired gut homeostasis, potentially impacting host health.

We next focused on the 10C group because it reflects a moderate daily intake of DPCs. Compared to the 0C control, the 10C group showed significant changes in the Firmicutes phylum (Supplementary Fig. 7c), particularly in probiotics-related genera. Enriched genera included *Parabacteroides* (Supplementary Fig. 7d), known for immune regulation and inflammation mitigation (*39*), *Oscillibacter*, involved in glucose metabolism (*40*), and *Dysosmobacter*, which may help alleviate obesity (*41*). Depleted genera such as *Turicibacter* (Supplementary Fig. 7d) are associated with anti-obesity and anti-inflammatory effects (*42*); *Ligilactobacillus* are important for stress regulation and pathogen defense (*43*). These findings suggest moderate MPs intake may significantly disturb cecal microbiota functions related to metabolism and immune systems.

To further explore the alterations in microbiome functions, we applied RSA to the cecal microbiota’s functional reaction pathways (RXN) and metabolic pathways (PWY) using the MetCyc database. We observed dose-dependent increases in biosynthetic pathways, particularly fatty acid biosynthesis. In contrast, pathways related to degradation of D-galactarate, citrate, myo inositol, and purine nucleobases, showed decreased abundance (Fig. 4e, f). The phyla Proteobacteria, Firmicutes, and Bacteroidetes were the main contributors to these functional shifts (Supplemental Fig. 7e). Previous research has correlated MPs exposure with microbiota changes that promote obesity (*44*) and associated metabolic disorders (*21*, *44*). Our findings align with these studies, suggesting impaired metabolic health (Fig. 2c).

We again focused on the functional differences between the 10C and 0C groups; distinct metabolic shifts were observed (Supplementary Fig. 7f). For example, the most pronounced increases were related to purine catabolism. Fatty acid biosynthesis pathways also showed marked upregulation. Conversely, the largest decrease was in the pathways degrading chlorinated phenols, implying a diminished capacity to detoxify these environmental pollutants in the presence of MPs. On the other hand, severe downregulations were noted in pathways related to the biosynthesis of amino acids, nucleotides, and energy metabolism. The sharp decline in tRNA charging and de novo biosynthesis pathways for adenosine and guanosine deoxyribonucleotides indicates a substantial systematic downshift in the synthesis of proteins and DNA nucleotides, respectively.

### MPs exposure led to tissue-specific transcriptomic aberrations with strong health implications

We delved into the impacts of MP exposure on gene expression across blood, placental, and mammary gland tissues. Using RNA sequencing and the Kallisto-sleuth pipeline (*45*), we observed tissue-specific expression profiles (Supplemental Fig. 8a), with blood and placental tissues exhibiting divergent overall expression patterns after exposure (Supplemental Fig. 8b-d).

Differentially expressed genes in response to varying MP exposure (1C, 10C, and 100C) were found in all three tissues (Supplementary Fig. 9a-c). Specifically, 796, 832, and 248 genes from blood, placenta, and mammary gland showed dose-dependent variations, forming two clusters via GLM (*p*-value < 0.01; Supplementary Fig. 9d). This led us to further explore dose-responsive functional dynamics. RSA pathway analysis, based on KO-KEGG pathway relationships, revealed dose-dependent impact in 142, 153, and 183 KEGG pathways in the blood, placenta, and mammary gland, respectively (Fig. 5a). The shared pathways between blood and placenta reflect physiological adaptations during pregnancy, including immune, hormonal, and environmental responses (*46–48*).

**Fig. 5.**
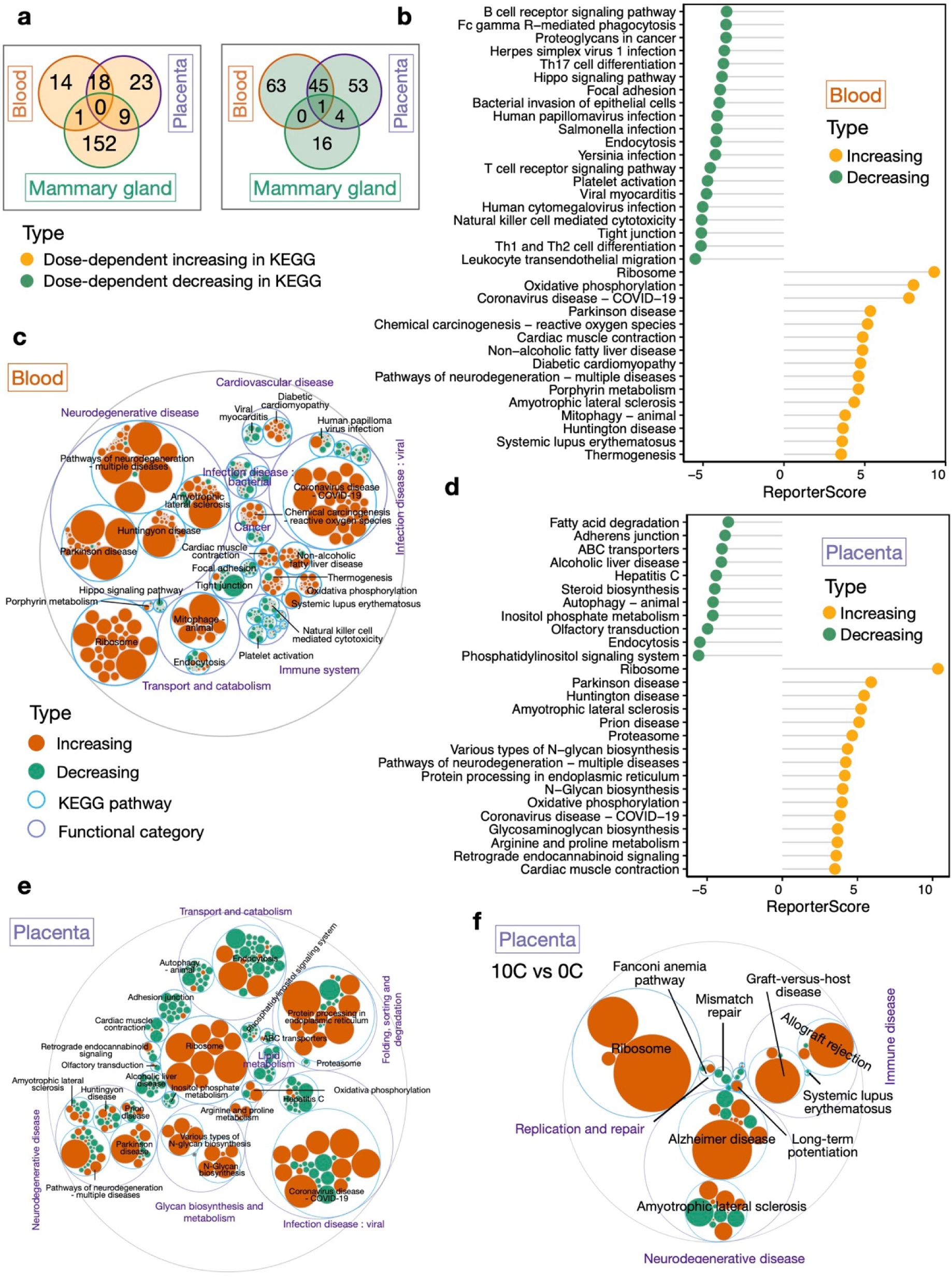
Transcriptomic functional changes in blood, placenta, and mammary gland in response to MPs exposure. (a) Venn diagram depicting pathways with significant (|ReporterScore| > 1.64, p < 0.05) increases (yellow) or decreases (green) in maternal blood, placenta, and mammary gland. (b) Lollipop charts displaying pathways with significant (|ReporterScore| > 3.5, p.adj < 0.01) increases (yellow) or decreases (green) in blood. (c) Gene-function integration maptree for blood, based on (b), showcasing functional categories, KEGG pathways, and gene levels, with the size of filled circles representing average gene expression abundance in four groups and their dynamic trends (increasing in red, decreasing in green). (d) Lollipop charts displaying pathways with significant (|ReporterScore| > 3.5, p.adj < 0.01) increases (yellow) or decreases (green) in placenta. (e) Gene-function integration maptree for placenta, based on (d), showcasing functional categories, KEGG pathways, and gene levels, with the size of filled circles representing average gene expression abundance in four groups and their dynamic trends (increasing in red, decreasing in green). (f) Gene-function integration maptree for placenta related to differential enriched pathways between 10C and 0C groups (|ReporterScore| > 2.0, p < 0.05), showcasing functional categories, KEGG pathways, and gene levels, with the size of filled circles representing average gene expression abundance in 10C and 0C groups and their differential trends (red for enriched in 10C; green for depleted in 10C).

In maternal blood samples, we observed dose-dependent increases in pathways associated with neurodegenerative diseases, viral infections, and ribosomal function, potentially leading to increased oxidative phosphorylation and oxidative stress (Fig. 5b, c, and Supplementary Fig. 10a, b). A decrease in cell adhesion pathways and an upsurge in mitophagy suggest potential protective mechanisms against oxidative damage (Fig. 5b, c). The blood samples also showed increased pathways related to cardiac muscle contraction, reflecting maternal cardiovascular adaptations (Fig. 5b, c). Comparing the 10C group with the 0C control group, we identified significant functional changes. The most enriched KEGG pathways in the 10C group were associated with infection by *Staphylococcus aureus* (Supplementary Fig. 10c, d), a critical human pathogen responsible for a broad spectrum of clinical infections. Additionally, pathways of neuroactive ligand-receptor interaction and glutathione metabolism were also upregulated. In contrast, the most depleted KEGG pathways in the 10C group included endocytosis, the hippo signaling pathway, and the degradation of branched-chain amino acids such as valine, leucine, and isoleucine (Supplementary Fig. 10c, d)

In the placenta data, we again observed dose-dependent increases in pathways related to neurodegenerative diseases, viral infections, ribosomal function, and protein processing in the ER, largely mirroring the changes in maternal blood (Fig. 5d, e, and Supplementary Fig. 11a, b). A decrease in adhesion junction pathways and autophagy within the placenta was noted, which may have implications for fetal development and pregnancy health (Fig. 5d, e). In terms of placenta-specific changes, an upregulation in glycan biosynthesis pathways was observed (Fig. 5d, e and Supplementary Fig. 11b), why may implicate reproductive complications, including preeclampsia, intrauterine growth restriction, and an increased risk of miscarriage (*49–51*). Conversely, we observed a downregulation of pathways involved in lipid metabolism, phosphatidylinositol signaling, inositol phosphate metabolism, and ABC transporters (Fig. 5d, e), which may adversely affect fetal development and the overall health of the pregnancy (*52–54*). Significant functional differences between the 10C and the 0C groups were identified (Fig. 5f and Supplementary Fig. 11c). In the 10C group, the most enriched pathways were associated with neurodegenerative diseases, ribosomal function, and immune diseases, while pathways related to replication and repair were depleted. The enrichment of pathways such as allograft rejection and graft-versus-host disease within the immune disease category is of special interest and suggests increased risks of miscarriage (*55*).

The mammary gland showed dose-dependent upregulation of pathways associated with infections and cancer, suggesting an active immune response to MPs exposure or potential pathological changes (Supplementary Fig. 12a). Conversely, the downregulated pathways were involved in metabolism and energy production (Supplementary Fig. 12a, b) and may reflect a shift in the glandular functional state or a compensatory mechanism in response to the increased demands of other pathways in the organ.

In conclusion, MPs exposure during pregnancy can alter gene expression in maternal blood, placenta, and mammary gland in a tissue-specific manner. The comparison between the 10C and 0C groups suggests that realistic levels of mixture MPs exposure can negatively impact metabolic, immune, neuronal, and reproductive functions during pregnancy.

### Benchmark Dose Model analysis via gene biomarkers indicates a safe consumption limit of 2-4 DPCs daily

In toxicological and health risk assessments, determining the threshold for adverse effects is essential. We integrated physiological (Fig. 2) and tissue-specific biomarker endpoints (Fig. 7a), which showed significant dose-responsive changes, to calculate benchmark doses (BMDs) of DPCs. BMDs, the preferred method for risk assessment by US EPA and the European Food Safety Authority (*56*), represent doses that elicit a specific change in response to an adverse effect. We estimated the lower 95% confidence intervals (BMDL) of BMDs for various endpoints using the US EPA benchmark dose software (BMDS; Methods) (*57*).

Our findings indicate that while physiological endpoints typically had higher safety limits, ranging from 15 to 38 cups daily during pregnancy, cecal microbiome and gene expression data request a lower limit of 2 to 4 cups daily (Fig. 7b). Several tissue-specific biomarkers, associated with inflammation, neurodegenerative diseases, and fetal development, were dose-responsive to MPs exposure. Notably, there was an increase in *Alistipes onderdonkii* and *Alistipes communis* in the cecum, linked to inflammation and cancer (*36*). In blood, placenta, and mammary gland, gene biomarkers such as *Ptk2b*, *Uba52*, *Hbq1a*, *Anxa5*, *Ddx19b*, *Pycr1*, *Nqo2*, *Clec5a*, and *Rpl26-ps6* were included, suggesting risks associated with impaired neuronal activity, DNA damage repair, compromised placental homeostasis, and increased oxidative damage and inflammation (*58–65*). These observations highlight the heightened sensitivity of including molecular-level endpoints in BMDs assessments.

In conclusion, moderate MP consumption of 3.3 cups/day during pregnancy could potentially disrupt gene expression across various tissues during pregnancy. Considering the varied sources of MPs intake, even non-DPC users reach the 2 to 4-cup exposure level.

**Fig. 6.**
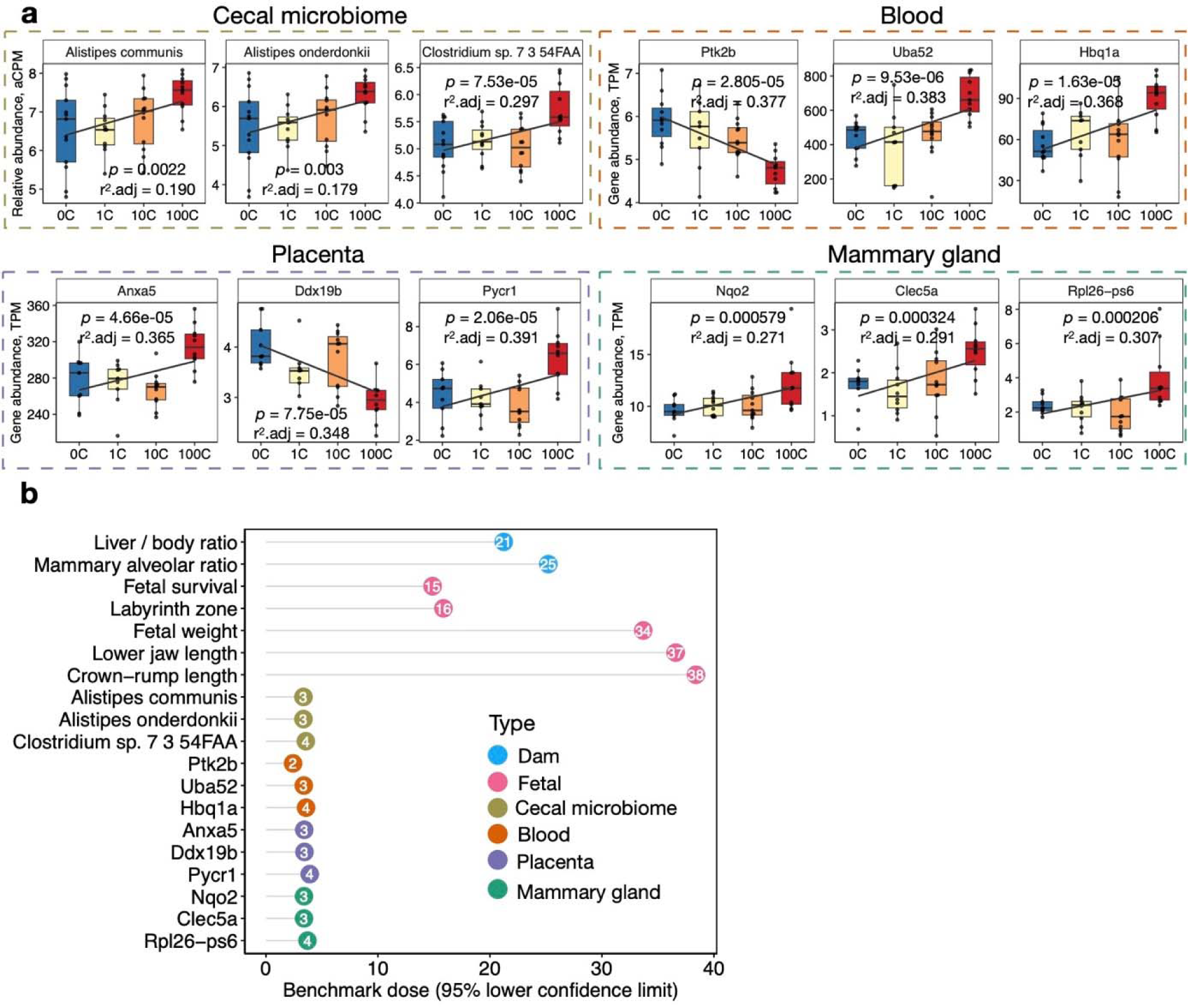
BMDL of MPs from DPCs in daily consumption. (a) Identification of top tissue-specific biomarkers, annotated with p-values and adjusted r-squared (R².adj) values derived from GLM. (b) Lollipop chart displaying the BMDLs evaluated using maternal physiology, fetal growth indices, and top tissue-specific biomarkers identified in (a).

## Discussion

The ubiquity of MPs and their potential health risks have become a focal point of environmental and public health research. Our heavy reliance on DPCs, including those made from plastic and plastic-coated paper, poses a significant source of MPs pollution, especially when these containers are used for hot beverages. While prior research has shed light on the release of MPs from these products (*25*, *26*), the full extent of their impact—from release to bioaccumulation and their toxicological effects at realistic exposure levels—remains underexplored. Existing research that investigated the health impact of MPs often used fixed-size plastic particles to simulate exposures, overlooking the size and type variations of MPs in real-world conditions, administered MPs by incorporating them into drinking water, leading to imprecise MPs intake (*21*, *23*), hindering the extrapolation to human-relevant dosage. Additionally, traditional detection methods, such as stereomicroscopy, SEM, FTIR, Raman, and fluorescent tagging with Nile Red using fluorescence microscopy, cannot capture the comprehensive size distribution of MPs in environmental or biological samples (*10*, *14*).

Our study aims to evaluate the full complexity and impact of MPs originating from DPCs. The MPs exposure to mice was calibrated to reasonable human exposure levels at 0.3/3.3 cups per day, making the results more generalizable. Our data reveal systematic dose-dependent deleterious effects on fetal development and physiological health of the dams. MPs were detected in all 13 examined tissues in a dose-responsive manner. The highest density of MPs was in cecal contents, but significant depositions were found in the fetus, placenta, kidney, spleen, lung, and heart. We propose a realistic MPs bioaccumulation model based on the quantity and size distributions observed in different tissues. In this model, MPs are absorbed from the gastrointestinal tract into the bloodstream and distributed to various organs (Fig. 3 b-d). Consequently, blood displayed the lowest MPs count but with a wide variation in size. Organs requiring a large amount of blood flow during pregnancy are embedded with a large number of capillaries with thin cell walls, such as placenta, kidney, spleen, and lung. These organs accumulated the majority of MPs. As expected, the BBB of brain would sift the majority of MPs in terms of both size and quantity, but some still got through. Importantly, the quantity and size of MPs did not differ significantly between placenta and fetuses, reflecting the unique physiological role of placenta in transferring a large number of large and small biochemical molecules, including MPs, to the developing fetus.

Mechanistically, we observed dose-responsive alterations in the cecal microbiota composition and function, particularly an increase in fatty acid biosynthesis pathways. Similar disturbances in the transcriptome were observed in maternal blood, placenta, and mammary gland in a tissue-specific manner. Importantly, even at 3.3 cups per day (the 10C group), we observed significant changes in cecal microbiome composition and metabolic functions, as well as gene expression profiles in maternal blood, placenta, and mammary glands. These changes were identified in several essential biological pathways, potentially affecting metabolic and immune functions, and increasing the risk of neurodegenerative diseases and miscarriage. The BMD analysis suggests that the safety margin for MPs exposure to DPCs during pregnancy lies between 2 to 4 cups daily, based on molecular-level endpoints. It is important to note that while previous research indicates that consuming 2-3 cups of coffee daily can benefit human health, they overlooked the potential release of MPs from DPCs (*28*).

Our study had several limitations. The detection of MPs was restricted to particles larger than 6 μm, and we quantified MPs based on particle counts, not mass. The in-depth characteristics of MPs in different tissues have not been determined. Our BMD results were based on molecular endpoints, which are more sensitive. Future studies should focus on quantifying the daily cumulative mass of MPs from all sources to better assess the physiological and molecular toxicity of realistic MPs exposure.

In summary, our study provided a more detailed understanding of the physiological and molecular consequences of exposure to MP mixtures at realistic levels in biological systems, particularly in the context of maternal and fetal health. Importantly, the 3.3 cups/day consumption is well within the range of the general population and the safety margin assessed by BMD, hence the adverse impact of MPs exposure on fetal development, cecal microbiome, immune health, neural systems, metabolic and reproductive functions are potentially generalizable to human health. This research calls for potential revisions in food safety standards, particularly considering vulnerable populations such as pregnant women and fetuses.

## Materials and methods

### Selection of coffee paper cups and MP sample preparation

For this study, we selected unused 500 ml DPCs from five popular brands, designated A through E. MP samples were then prepared using three cups from each brand. Following the procedure outlined by a prior study (*14*), we poured 420 ml of hot ultrapure water (filtered by 0.22 μm filter, temperature 85–90°C, pH∼6.9) into each cup and let it stand for 15 minutes. This time frame corresponds with consumer habits, where most report finishing their beverages within this duration (*14*).

### MPs detection and quantification

To ascertain and quantify MPs, we sampled the mixed hot ultrapure water from each cup. MPs were identified using Nile Red, a lipophilic dye that binds to plastics, causing them to fluoresce under specific light conditions (excitation: 534–558 nm, emission: > 590 nm). From each sample, 10 μL was smeared on a slide, stained with 0.01 mg/ml Nile Red, and oven-dried at 55 °C for 30 minutes. Fluorescence microscopy (Nikon, Tokyo, Japan) was used for visualization. Controls included cups with room-temperature water (Supplementary Fig. 1). All experiments were done in triplicate.

### Preparation of concentrated MP solution

We selected the brand with the highest MP concentration to prepare a concentrated MP solution. MPs were extracted from eighty disposable paper coffee cups via heat condensation at 85 °C, and their concentration was assessed as described before. We used a dose conversion factor to translate human doses to mouse equivalents (*66*): Animal equivalent dose (cups/kg) = Human dose (cups/kg) × 12.3 Given an average human weight of 60 kg and a mouse weight of 0.035 kg, a single human cup translates to 0.0071715 cups for mice. We thus prepared MP solutions corresponding to 1, 10, and 100 human cups, resulting in mouse doses of 0.007175, 0.07175, and 0.7175 cups, respectively, in a 200 µl volume.

### Animal experiments

The animal experiments followed Zhejiang University’s Institutional Animal Care and Use Committee protocols (Approval number: ZJU20230176). Eight-week-old specific-pathogen-free C57BL/6J mice, sourced from the university’s Laboratory Animal Center, were maintained under optimal conditions, including a temperature of 21 ± 3°C, humidity maintained at 50%, and a 12-hour light-dark cycle. Nine-week-old female mice (approximately 20 g), after a week of acclimatization, were coupled with males at a ratio of 3:1, with pregnancy confirmed by seminal plug presence, defining gestational day 0 (GD0) (*67–69*). Post-mating, males were removed, and females were allocated into groups, standardized by body weight to start the experiment uniformly.

On gestational days 3, 6, 9, 12, and 15, control group (0C) mice were gavaged with 200 μL of ultrapure water. Mice in the experimental groups received 200 μL of a MP solution, corresponding to the human equivalent of consuming 1, 10, or 100 coffee cups, named as 1C, 10C, and 100C, respectively. On day 18, mice were euthanized with CO_2_, followed by cervical dislocation. Maternal blood was collected from the posterior orbital venous plexus and stored at −80°C for subsequent transcriptomics analysis. The first leftmost placenta was fixed in 10% neutral buffered formalin and hematoxylin and eosin (HE) stained, and the first rightmost placenta was snap-frozen in liquid nitrogen for transcriptomic analysis.

The weight and survival status of fetuses from each dam were meticulously recorded. For morphometric analysis, fetuses were photographed next to a calibrated ruler, and measurements of the lower jaw and crown-rump lengths were determined using Fiji image analysis software. The histological examination involved the processing of the left fourth inguinal mammary gland with H&E staining, while the right fourth inguinal mammary gland was snap-frozen in liquid nitrogen for transcriptomic analysis. Cecal contents were also collected, immediately frozen in liquid nitrogen, and stored at −80°C for subsequent metagenomic sequencing.

In addition to the reproductive endpoints, 13 types of tissues were systematically collected, including the cerebral cortex, hippocampus, cerebellum, heart, liver, spleen, lung, kidney, the right fifth inguinal mammary gland, the first fetus from the leftmost uterine horn, and the second placenta from the left side of the cervix. These tissues were weighed, snap-frozen in liquid nitrogen, and preserved at −80°C to assess MP size distribution and accumulation. The overall experimental procedure, from collection to analysis, is depicted in Fig. 2a.

### Tissue sample digestion

Tissue digestion was conducted in two stages, adapting established methodologies with minor modifications (*12*, *13*, *70*). To prepare the digestion solution, a 10% w/v potassium hydroxide (KOH) solution was made using ultrapure water passed through a 1.6 μm filter membrane. KOH pellets, procured from Sigma-Aldrich, were dissolved in ultrapure water to achieve the desired concentration. Each tissue specimen was placed in an individual glass container, to which 10 ml of the KOH solution was added. The containers were securely sealed to prevent evaporation and contamination and then placed in a constant-temperature oven set to 60°C for a digestion period of seven days.

Following initial digestion, 5 ml aliquots of the sample suspension were vacuum-filtered through a 0.45 μm-pore-size glass fiber filter (47 mm diameter; Shanghai Dibo Biotechnology, China). This filtration process was conducted within a laminar flow hood to maintain a sterile environment. To defat the samples, each aliquot was washed with 10 ml of acetone, followed by a rinse with 10 ml of ultrapure water to facilitate resuspension and minimize premature filter clogging. The filters were then returned to glass containers for the second digestion phase. Here, 10 ml aliquots of 30% hydrogen peroxide (H_2_O_2_) were added twice at 24-hour intervals. After an additional 48 hours to allow for the attenuation of H_2_O_2_’s oxidative potential (*70*), the samples underwent ultrasonic treatment at 60 kHz for 30 minutes. The resulting suspensions were then rendered suitable for subsequent analysis.

MP extraction from cecal contents was performed with the method outlined in a previous study (*71*). In summary, Fenton’s reagent, which facilitates organic matter degradation, was freshly prepared by combining 30% hydrogen peroxide (H_2_O_2_) with an iron catalyst solution. The latter was made by dissolving 20 g of iron (II) sulfate heptahydrate in 1 L of ultrapure water. These two components were then mixed in glass beakers, using a volume ratio of 1 part iron solution to 2.5 parts H_2_O_2_. Cecal contents were then treated with 20 mL of this Fenton solution and allowed to react at room temperature for 5 hours.

Following the Fenton reaction, the mixture was filtered through a cellulose nitrate– cellulose acetate filter (CN-CA filter; Shanghai Dibo Biotechnology). To ensure complete digestion of organic matter, the filters were subsequently treated with 10 mL of 65% nitric acid (HNO_3_) at 50°C for 30 minutes, and then the temperature was increased to 70°C for an additional 10 minutes. The digested mixture was then diluted with ultrapure water in a 1:2 volume ratio. This final solution is ready for analysis, with the MP content made accessible for quantification and characterization.

To mitigate the risk of plastic contamination, all personnel were required to wear cotton lab coats, face masks, and disposable latex gloves while during all experimental procedures. Prior to any experiments, all surfaces were thoroughly sanitized using 70% ethanol. Furthermore, all aqueous solutions, including ultrapure water for cleaning and the preparation of the KOH solution, were filtered through a 0.45 μm-pore-size membrane (Whatman GF/A) to remove any particulates. Glassware and tools, such as filter funnels, scissors, tweezers, and scalpels, underwent a rigorous cleaning protocol. This involved washing with dish soap, a preliminary rinse with acetone, soaking in ultrapure water, and a final rinse with ultrapure water that had been filtered through a 0.45 μm membrane, ensuring the removal of potential contaminants. To account for and adjust for any inadvertent contamination, three procedural blanks were prepared following the identical protocol used for the samples, but without the inclusion of any tissue. These blanks were maintained in close proximity to the actual samples throughout the experimental process to serve as controls for assessing background contamination levels.

### Raman Microspectroscopy analysis

For the Raman Microspectroscopy analysis, 2 mL of the digested tissue sample was filtered through a 0.45 μm-pore-size glass fiber filter (47 mm diameter; Shanghai Dibo Biotechnology) using a vacuum pump attached to a filter funnel, all within a laminar flow hood to maintain a contaminant-free environment. After filtration, the filter was washed with 10 mL of ultrapure water and left to air-dry at ambient room temperature. Once dry, the filters were stored in glass Petri dishes for subsequent analysis.

The dried filter membranes were then analyzed using a Raman XploRA Nano Microspectrometer (Horiba Scientific), which is equipped with a 785 nm laser diode and a grating of 600 lines per mm, providing a spectral range of 0-2200 cm^-1^. Calibration of the instrument was performed using the silicon line at 520.7 cm^-1^ to ensure accurate spectral data.

To process the raw Raman spectra, polynomial baseline correction and vector normalization were applied using Labspec 6 software (Horiba Scientific). These steps were crucial for reducing noise and improving the clarity of the spectral data. MP identification was conducted by comparing the obtained spectra against reference spectra in the SLOPP Library of Microplastics and the KnowItAll software’s spectral library (Bio-Rad Laboratories, Inc.). Matches achieving a Hit Quality Index (HQI) score of 80 or above were considered to be reliable identifications of the MPs.

Additionally, 10 μL of a concentrated MP solution from the group designated for gavage (100C group) was placed onto a filter and analyzed using the same Raman microspectroscopy methodology. To ensure the reproducibility and reliability of the results, all analytical procedures were performed in triplicate.

### Analysis of MPs size distribution and accumulation

To assess MPs size distribution and accumulation, 10 μL of the homogeneously mixed tissue digest was combined with 190 μL of a 0.01 mg/mL Nile Red solution in a 96-well plate. The samples were incubated for 30 minutes at 55 °C to ensure proper staining of the MPs. Each tissue sample was processed in triplicate to ensure consistency, and each 96-well plate included three processing blank controls to account for background fluorescence.

The fluorescence of the stained MPs was measured using a BioTek Cytation 3 plate reader (Bio Tek Instruments, Inc., USA) at 4x magnification with three technical replicates. The Montage function of the plate reader was utilized to quantify MPs across the entire well. This approach enabled the accurate analysis of MPs as small as 6 μm in diameter. To avoid systematic errors, the same acquisition and analysis parameters were applied across all wells. For clarity in the resulting data, MPs were labeled for enhanced visualization in Fig. 1b.

Quantification of MPs accumulation was expressed as the total count of MPs per gram of tissue. For example, in the case of fetal tissue analysis, the fluorescence-derived count was multiplied by 1,000 (reflecting the total digestion volume of 10 mL) and then divided by the weight of the fetal tissue to normalize the results.

Furthermore, for evaluating MPs from the 0C, 1C, 10C, and 100C groups designated for gavage, 5 μL of a concentrated MP suspension was incubated with 195 μL of the Nile Red solution at a concentration of 0.01 mg/mL. This step was carried out in six replicates to ensure statistical robustness, following the same staining, incubation, and analysis protocol as previously described.

### DNA extraction, library preparation, and metagenomic sequencing

Genomic DNA was isolated from the cecal contents using the magnetic-based soil and stool DNA extraction kit (TIANGEN®, Beijing, China). The concentration of the extracted DNA was determined using the Qubit Fluorometric Quantitation system (Thermo Fisher Scientific, Waltham, USA).

For DNA library construction, we utilized the VAHTS Universal Plus DNA Library Prep Kit for Illumina® (Vazyme #ND617, Nanjing, China), following the kit’s protocol. This process included using adaptors and primers from the VAHTSTM Multiplex Oligos Set 5 for Illumina® (Vazyme #N322).

The prepared DNA libraries were then sequenced on an Illumina NovaSeq 6000 platform, utilizing a paired-end 150 bp (2×150 bp) configuration (Illumina Inc., San Diego, CA). The sequencing service was provided by Novogene Co., Ltd (Beijing, China).

### General statistical analysis and data visualization

We conducted our primary statistical analyses using custom scripts within a Linux environment and R (version 4.1.2) through RStudio. Data visualizations were also generated using RStudio, leveraging a suite of packages primarily sourced from the Bioconductor project (https://www.bioconductor.org/). The core packages utilized in our analyses included ggplot2 (version 3.3.5) for creating advanced graphics, ggbeeswarm (version 0.6.0) for bee swarm plots, gganatogram (version 1.1.1) for a mouse anatomical diagram, dplyr (version 1.0.8) for data manipulation, and ggpubr (version 0.4.0) for publication-ready plots. Additional packages that enhanced our visualizations were ggsci (version 2.9) for scientific journal-themed color palettes, ggrepel (version 0.9.1) for repelling overlapping text labels, and ggtreeExtra (version 1.4.1) for annotating phylogenetic trees. We also employed reshape2 (version 1.4.4) for data restructuring, RColorBrewer (version 1.1-2) for color palettes, vegan (version 2.5.7) for community ecology analyses, viridis (version 0.6.2) for colorblind-friendly palettes, circlize (version 0.4.15) for circular visualization, patchwork (version 1.1.2) for combining multiple plots, gtable (version 0.3.0) for arranging tables, grid (version 4.1.2) for grid graphics, and mvtnorm (version 1.1-3) for multivariate normal and t distributions. The R source code used for the analyses is available in the supplementary materials.

For the analysis of univariate variables such as fetal growth, non-parametric statistical tests (Kruskal-Wallis) were applied, unless the data conformed to a normal distribution, in which case parametric tests were used. For example, the pregnancy weight of dams was analyzed using a two-way ANOVA with an unbalanced design to account for variability. In addition, we utilized generalized linear models with logarithmic link functions to investigate dose-dependent trends in the data.

### Metagenomic data processing and statistical analysis

The taxonomic classification of the metagenomic sequencing reads was performed to characterize the comprehensive species composition within the cecal microbiota. We utilized the Kraken 2 (*31*) for this purpose, employing a custom-compiled database that included a wide range of genomic categories: bacteria (370,424 genomes), fungi (7,485 genomes), viruses (41,963 genomes), archaea (6,269 genomes), plants (Viridiplantae, 1,612 genomes), vertebrates (1,242 genomes), invertebrates (1,120 genomes), and protozoa (994 genomes). The figures in parentheses represent the number of genomes included for each taxonomic group. The comprehensive database resulted in a Kraken 2 pre-built index that was over 3 terabytes in size. The reference genomes for constructing this database were obtained from the RefSeq and GenBank repositories as of December 30, 2020.

For the normalization of our data, we employed the Counts Per Million (CPM) method, which was then transformed using the hyperbolic arcsine (arcsinh) function—specifically, the asinh function in R. All downstream statistical analyses and visualizations were based on these arcsinh-transformed CPM values (referred to as aCPM). We crafted annotated phylogenetic trees using custom Python scripts and the ggtreeExtra package in R.

To assess the differences in microbial community composition across our samples, known as β-diversity, we computed Bray–Curtis dissimilarity indices from the abundance data. We visualized the ordination using Principal Coordinate Analysis (PCoA) and conducted statistical tests to compare the communities using Analysis of Similarities (ANOSIM) and Permutational Multivariate Analysis of Variance (PERMANOVA), each with 999 permutations for robustness.

In addition, we utilized the HUMAnN3 (*72*) to detect shifts in gene expression from the clean metagenomic reads. These reads were aligned to the UniProt90 database (version 2019_01) to quantify gene expression at the reaction level and to the MetaCyc database for pathway-level expression. For taxonomic classification of the reads, MetPhlAn 3.0 (*72*) and the ChocoPhlan pangenome database were employed. Normalization of reads mapped to reactions and pathways was performed against sequencing coverage, and results were presented as copies per million (cpm, denoted as Copm) in the metagenomic datasets.

Functional enrichment analysis was performed using the ReporterScore package with a ‘directed’ model, available at https://github.com/Asa12138/ReporterScore.

### RNA extraction, library preparation, and RNA-sequencing

RNA was extracted from whole blood samples utilizing the PAXgene Blood RNA Kit. Concurrently, total RNA from placental and mammary tissues was isolated with the TRIZOL RNA Extraction Kit (Invitrogen, Carlsbad, CA, United States). The integrity of the extracted RNA was meticulously assessed through several quality control measures, including quantification with a Nanodrop spectrophotometer, evaluation for degradation and contamination using agarose gel electrophoresis, and comprehensive analysis of RNA integrity and concentration via the Agilent 2100 Bioanalyzer (Agilent Technologies, Santa Clara, CA, United States).

Library preparation for RNA sequencing was performed using the NEBNext Ultra RNA Library Prep Kit for Illumina (New England Biolabs, USA, Catalog #: E7530L), strictly adhering to the manufacturer’s protocol. Unique index codes were incorporated to enable sample identification after sequencing. The prepared libraries were sequenced on Illumina platforms by Novogene Co., Ltd.

### RNA-sequencing data processing and statistical analysis

The sequencing yielded 48.27 million base pair (bp) paired-end reads for blood, 33.75 million for placenta, and 38.46 million for mammary glands. For the quantification of transcript abundance, Kallisto (version 0.50.0) (*45*) was utilized to perform pseudoalignment. An index was constructed using the Mus musculus transcriptome fasta file from the GRCm39 build. To assess the inferential variance in transcript abundance, 100 bootstrap samples were generated. These reads displayed high mapping rates to the reference genome GRCm39, with 95.40% in blood, 94.50% in the placenta, and 92.83% in the mammary gland.

We conducted differential gene expression analysis using the sleuth (*73*), which integrates transcript-level p-values (*74*). The kallisto-sleuth pipeline was chosen for its enhanced sensitivity and more stringent control of the false discovery rate compared to alternative methods (*45*).

Pathway Annotation and Statistical Correlation Analysis involved annotating pathways based on the Kyoto Encyclopedia of Genes and Genomes (KEGG). This was executed within the ReporterScore package, employing the ‘directed’ mode.

### Benchmark dose calculation

To evaluate the toxicological responses in this study, we performed benchmark dose calculation using data showing significant dose-response relationships. This included both physiological and molecular markers, as illustrated in Fig. 7a. We employed a continuous endpoint approach for all features, with the exception of fetal survival, which was examined using a nested dichotomous endpoint framework.

The selection of the optimal BMD model within the US EPA BMDS framework was conducted through a consensus-based approach. This involved multiple criteria, including: a goodness-of-fit threshold with a p-value greater than 0.10; the lowest Akaike’s Information Criteria (AIC); scaled residuals less than the absolute value of 2; a BMD to BMDL (lower 95% confidence interval) ratio less than 5; and a visual inspection of the curve fit to ensure both plausibility and model parsimony (*57*).

For our analysis, the default confidence level was set at 95%, corresponding to a one-sided 95% confidence limit. We opted to use the lower confidence limit of the BMDL for deriving health guidance values, as it offers a more conservative and precautionary estimate of the toxic dose.

## Acknowledgments

This study was supported by the China Postdoctoral Science Foundation (2021M692823) and the Zhejiang University LSI start-up funding. We are grateful to our colleagues at the core facility of the Life Sciences Institute, especially the NECHO high-performance computing cluster.

## Author contributions

Q.C. and C.J. conceived the study. Q.C., C.P., Z. S. H. X., G. Y., and Y. S performed experiments. Q.C., C.P., and X.W. performed bioinformatic analyses. Q.C. and C.J. drafted and revised the manuscript with input from other authors.

## Data availability

The raw sequencing reads were submitted to ENA under project ID PRJEB70815.

## Additional information

Supplemental Fig 1-12.

