## Supplemental information for "Microplastics from disposable paper cups are enriched in the placenta and fetus, leading to metabolic and reproductive toxicity during pregnancy"

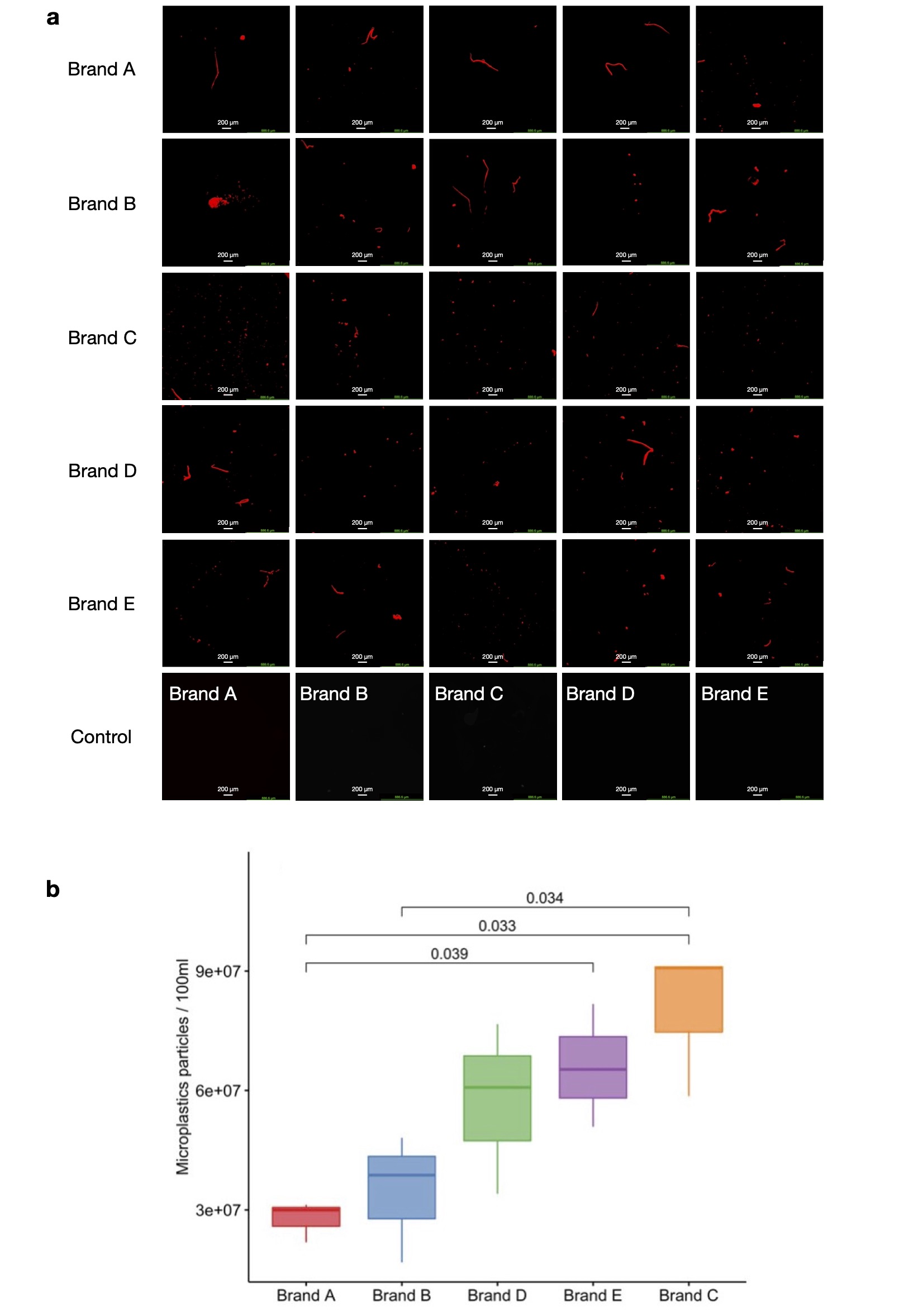

**Supplementary Fig. 1.** **Characteristics of microplastics released from disposable paper coffee cups of different brands.**

(a) Fluorescence microscopy images at 4X magnification showing microplastic particles in hot ultrapure water after a 15-minute incubation in paper coffee cups from Brands A to E. Control samples were treated with ultrapure water at room temperature for 15 minutes.

(b) Boxplots displaying the distribution of microplastic numbers from various brands, with statistical differences determined using the Kruskal-Wallis test and post hoc Games-Howell tests.

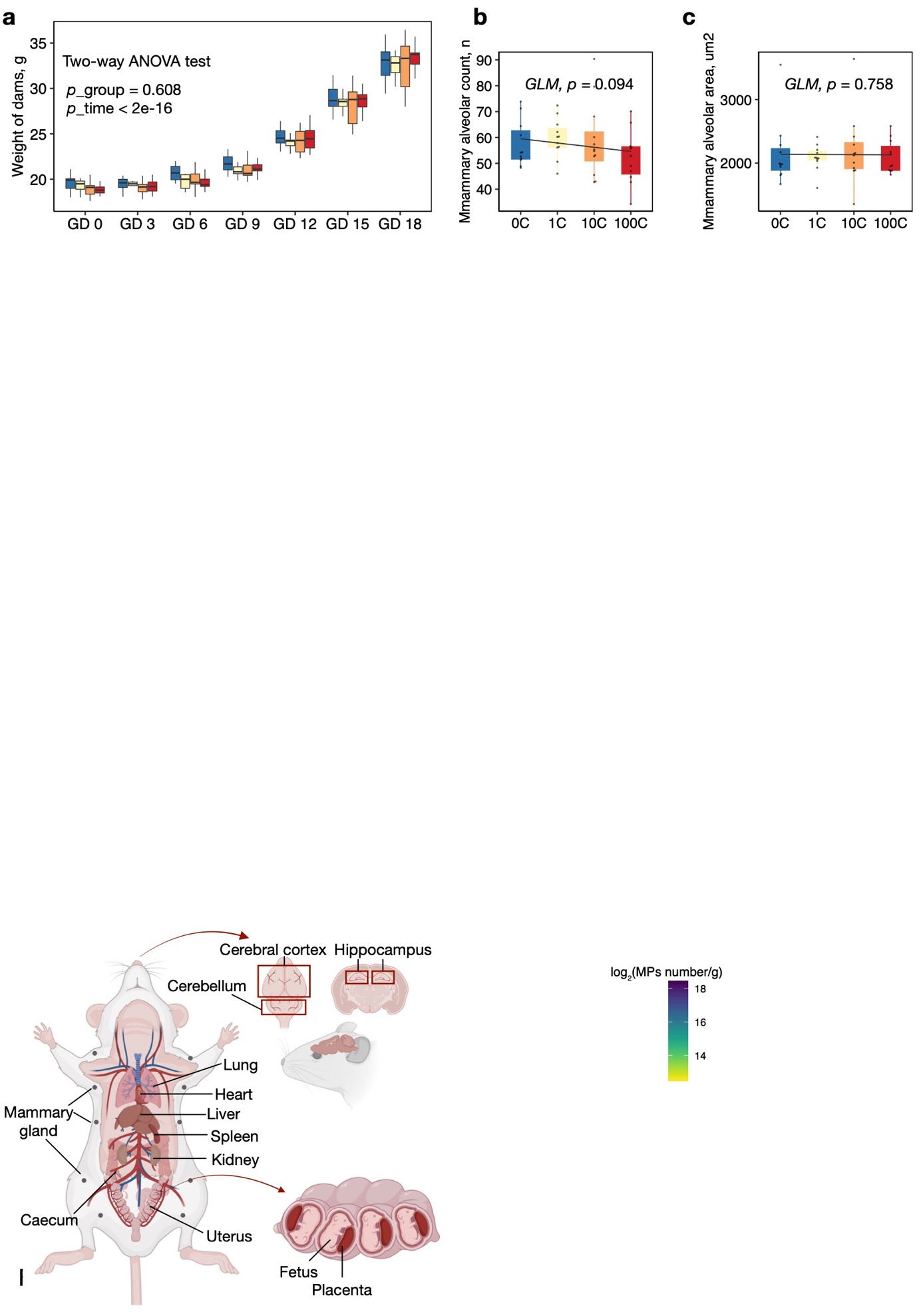

**Supplementary Fig. 2**. **Effects of microplastic exposure on maternal body weight, mammary alveolar count, and mammary alveolar area.**

(a) Maternal body weight variations.

(b) Mammary alveolar count variations.

(c) Mammary alveolar area measurements.

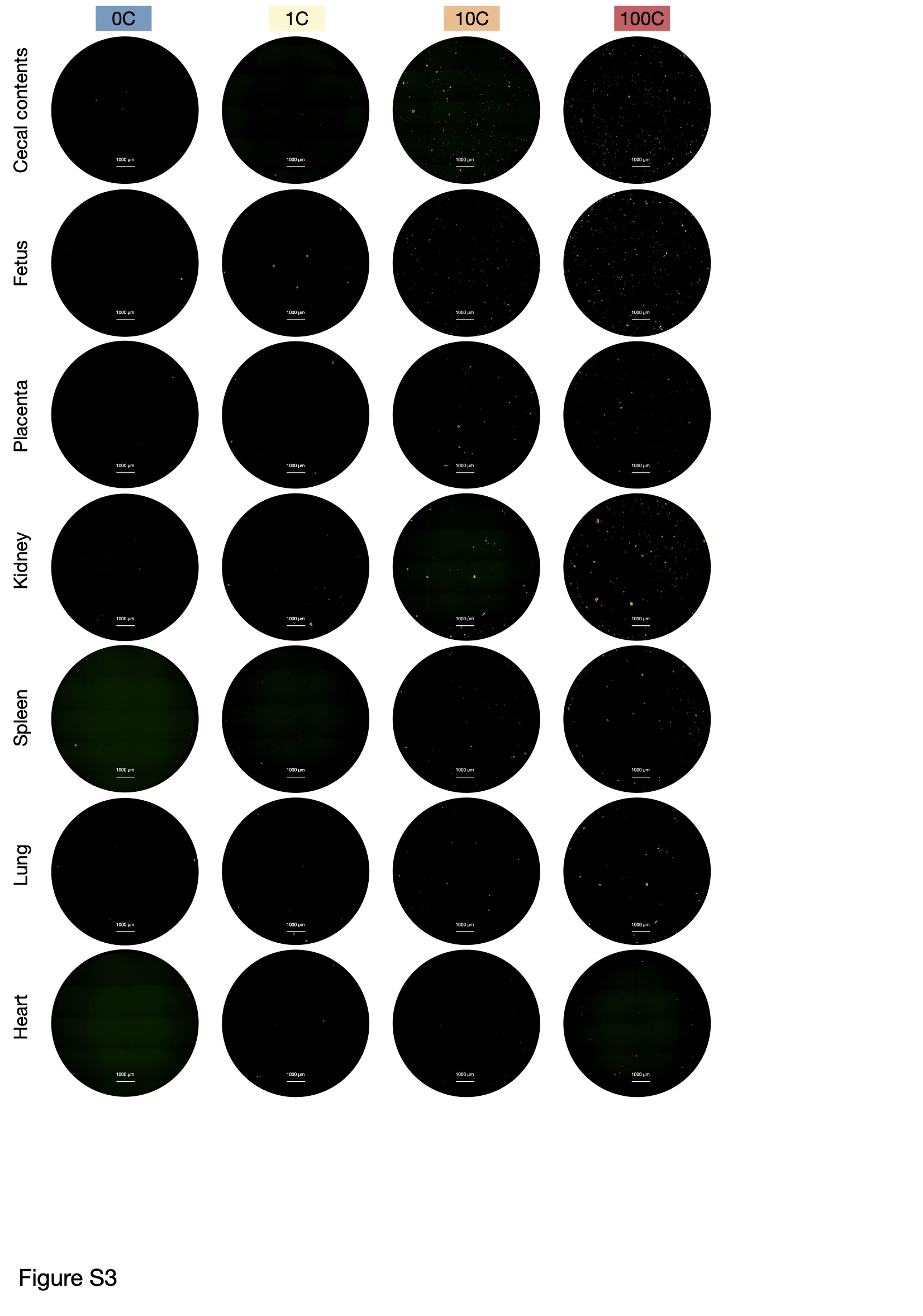

**Supplementary Fig. 3.** **Visualization of MPs in various tissues.** Representative images of cecal contents, fetus, placenta, kidney, spleen, lung, and heart from all groups. Each comprises 28 merged images at 4x magnification using the montage function.

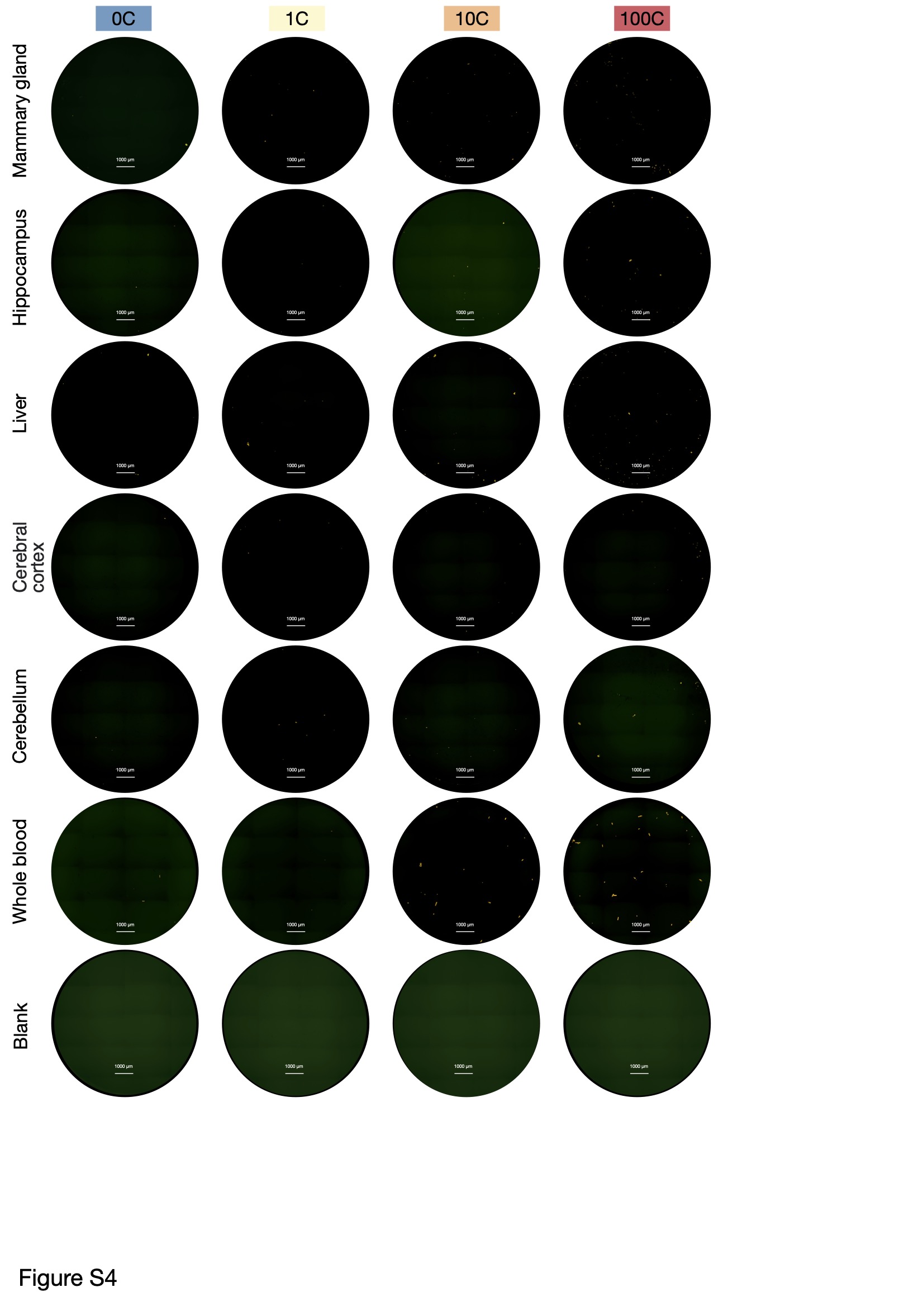

**Supplementary Fig. 4.** **Visualization of MPs in various tissues.** Representative images of the mammary gland, hippocampus, liver, cerebral cortex, cerebellum, whole blood, and blank control from all groups. Each comprises 28 merged images at 4x magnification using the montage function.

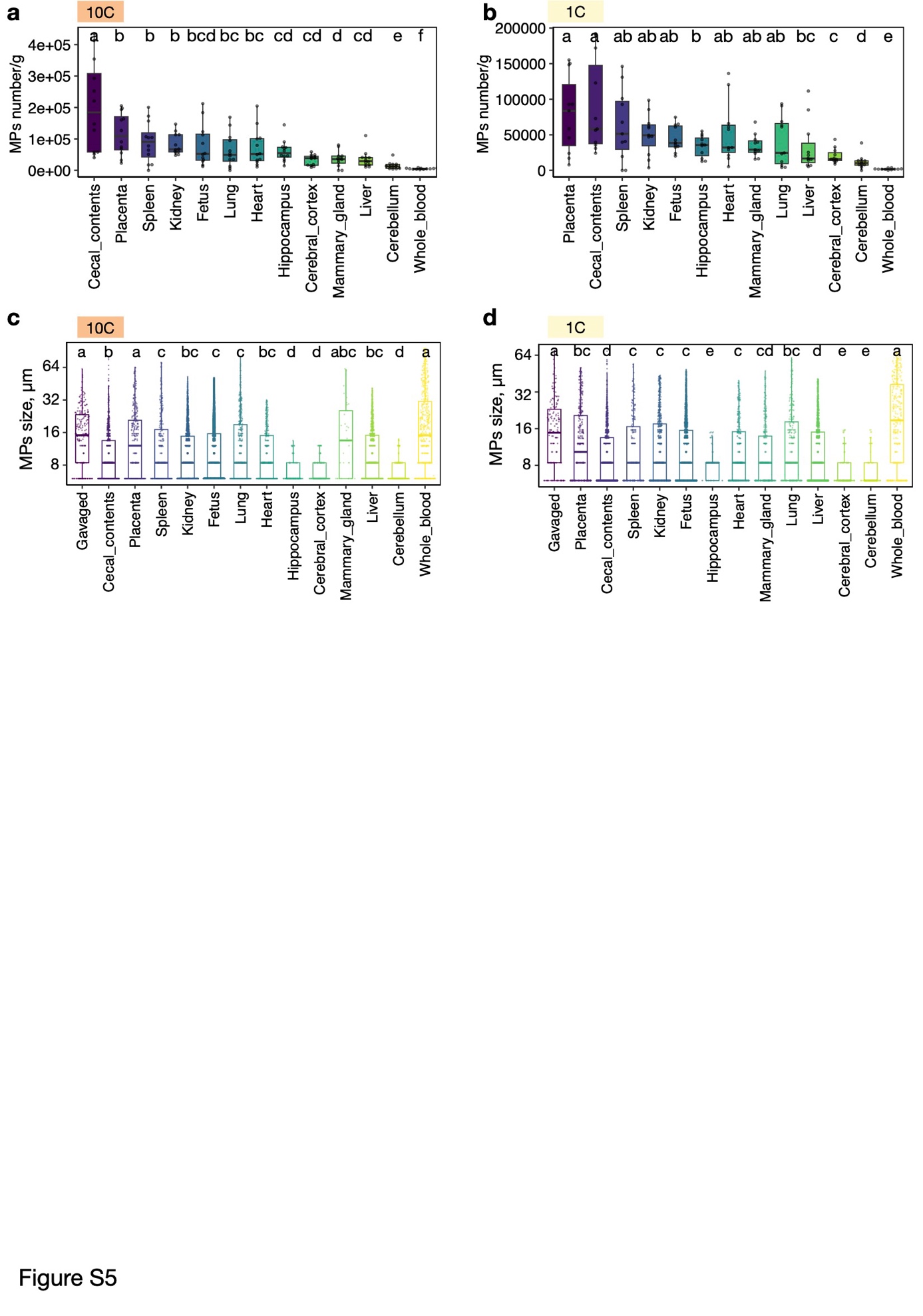

**Supplementary Fig. 5.** **Distribution and size variation of microplastics in biological samples of 10C and 1C groups.**

(a-b) Accumulation of MPs in different tissues in the 10C (a) and 1C (b) groups, with statistical significance denoted by lowercase letters for p < 0.05 via the Kruskal-Wallis test with post hoc Games-Howell tests.

(c-d) Size distribution of microplastics in the 10C (c) and 1C (d) groups, with statistical differences indicated by lowercase letters for p < 0.05, as determined by the Kruskal-Wallis test with post hoc Games-Howell tests.

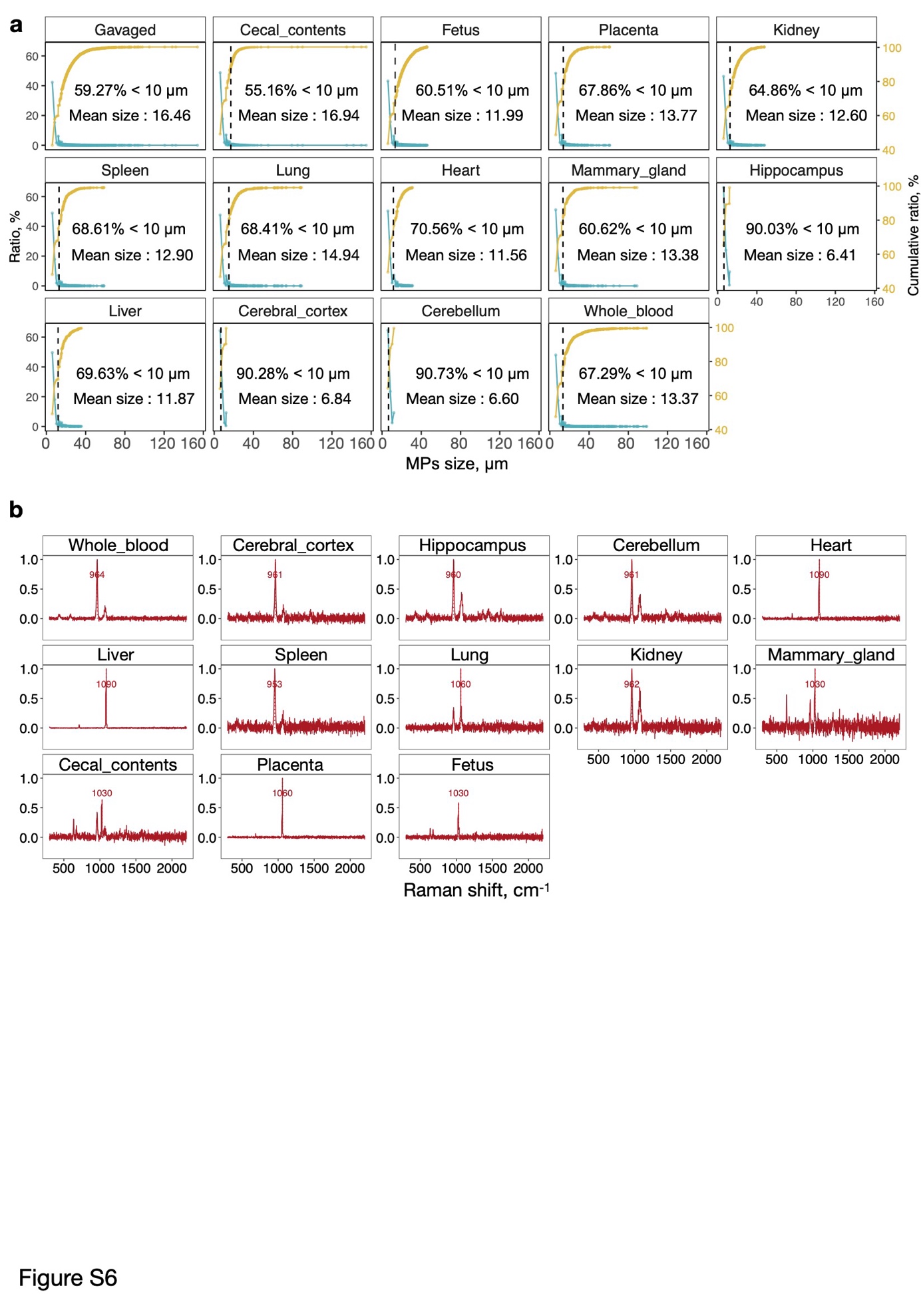

**Supplementary Fig. 6.** **Characteristics of microplastics in biological samples from the 100C group.**

(a) Frequency and cumulative distribution of particle sizes (blue for frequency, yellow for cumulative), with a vertical blank dashed line indicating the average particle sizes.

(b) Raman spectroscopy profiles of the microplastics.

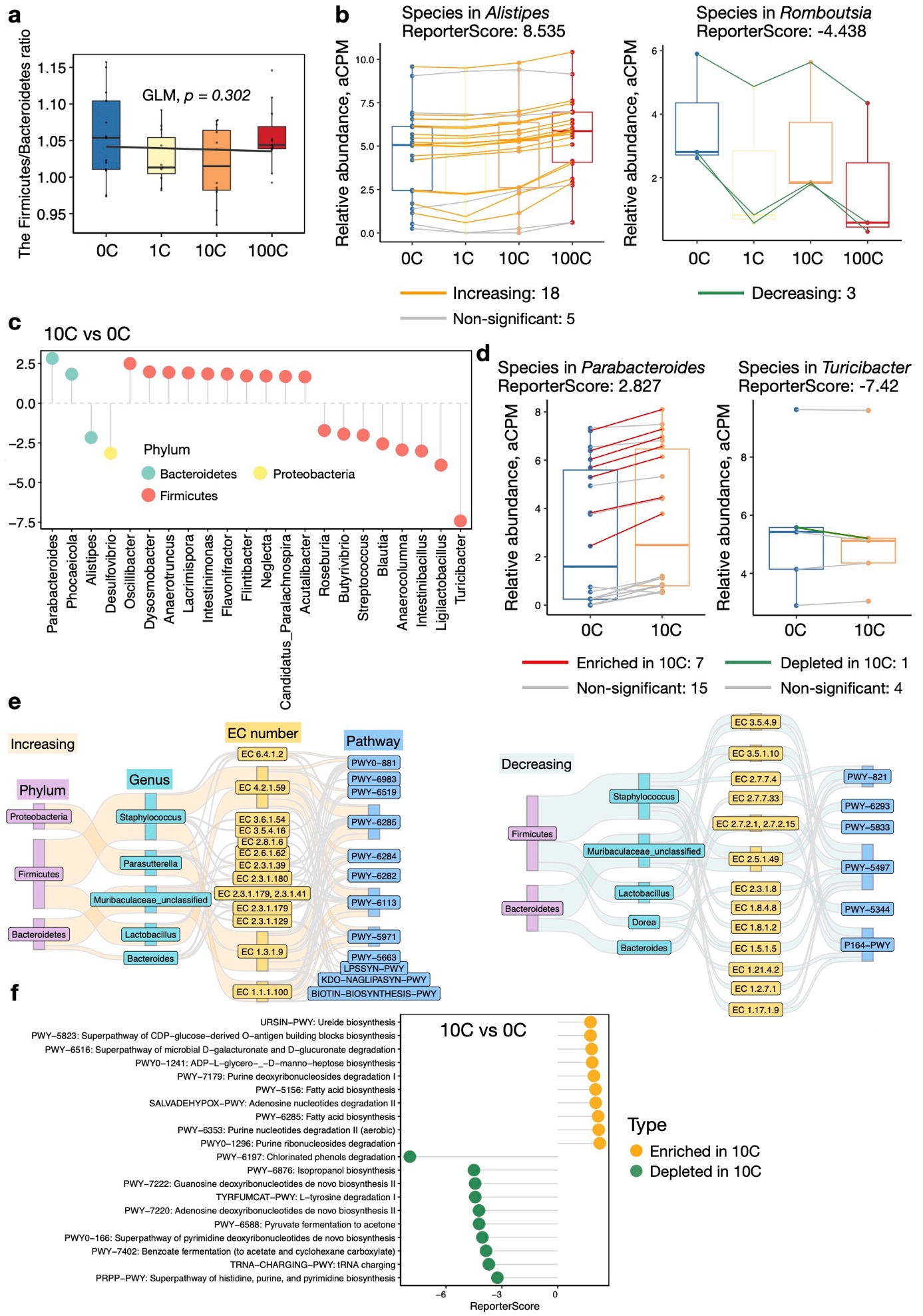

**Supplementary Fig. 7.** **Microbiome composition and functional enrichment analysis.**

(a) Firmicutes-to-Bacteroidetes ratio with GLM p-values.

(b) Box chart showing the abundance of genera *Alistipes* and *Romboutsia* across groups. Line colors indicate a trend in relative abundance, with *Alistipes* showing the largest positive and *Romboutsia* the largest negative reporter scores.

(c) Lollipop chart of significantly enriched genera (|ReporterScore| > 1.64, p < 0.05) in 0C and 10C groups.

(d) Box chart for genera *Parabacteroides* and Turicibacter in 0C and 10C groups, with line colors indicating abundance trends. *Parabacteroides* shows the largest positive and *Turicibacter* the largest negative reporter scores.

(e) Sankey diagrams depicting the associations between source taxa at the phylum and genus levels, EC numbers, and pathways, separated by increasing or decreasing ReporterScore in response to dose variations.

(f) Lollipop chart of significantly enriched PWYs (|ReporterScore| > 2.0, p < 0.05) in 0C and 10C groups.

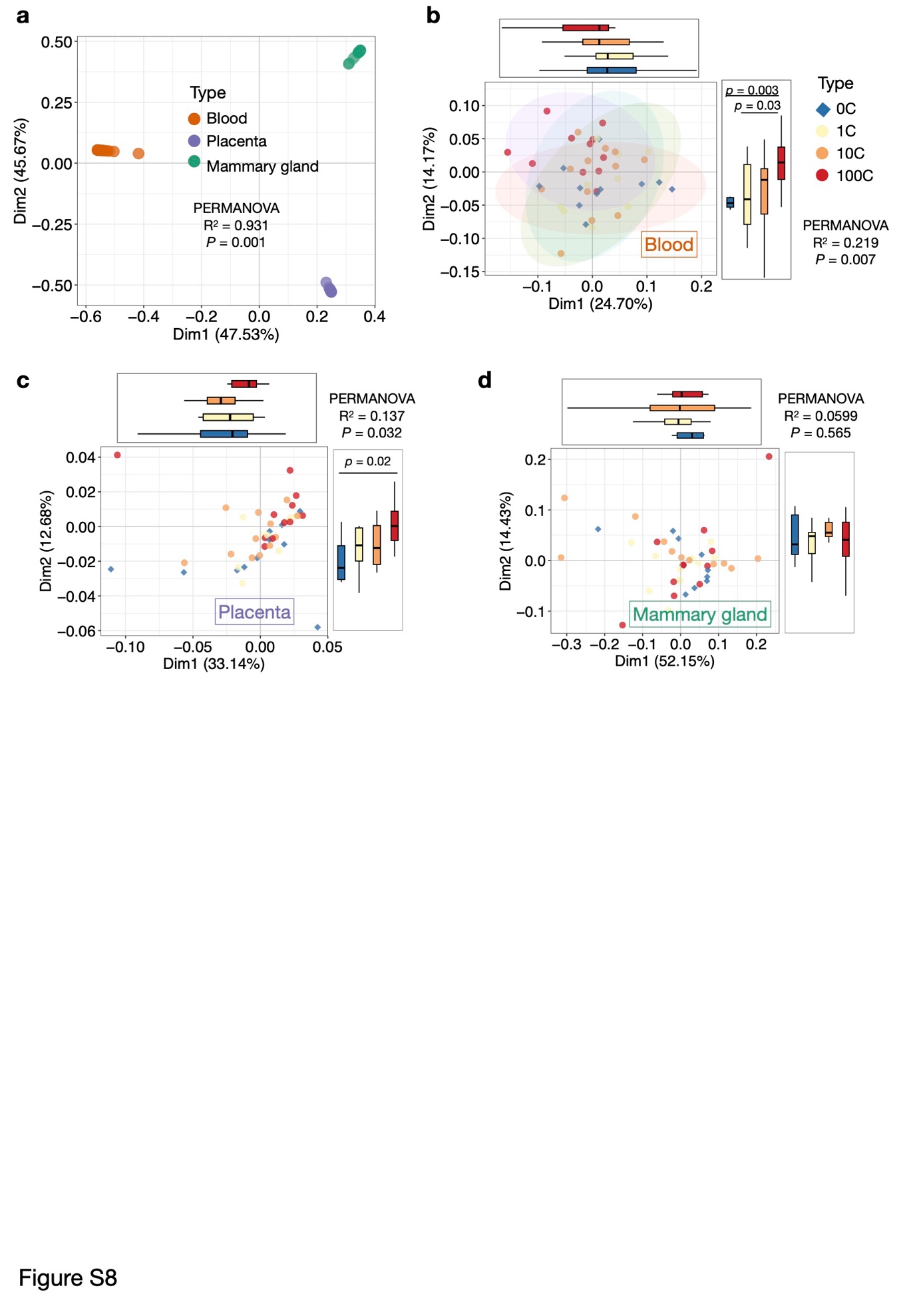

**Supplemental Fig. 8.** **Differential transcriptomic responses in blood, placenta, and mammary glands.**

(a) PCoA analysis of gene expression across tissues, with boxplot indicating statistical significance with Kruskal-Wallis test.

(b-d) PCoA analysis of gene expression across samples in the blood (b), placenta (c), and mammary gland (d), with boxplot indicating statistical significance with Kruskal-Wallis test.

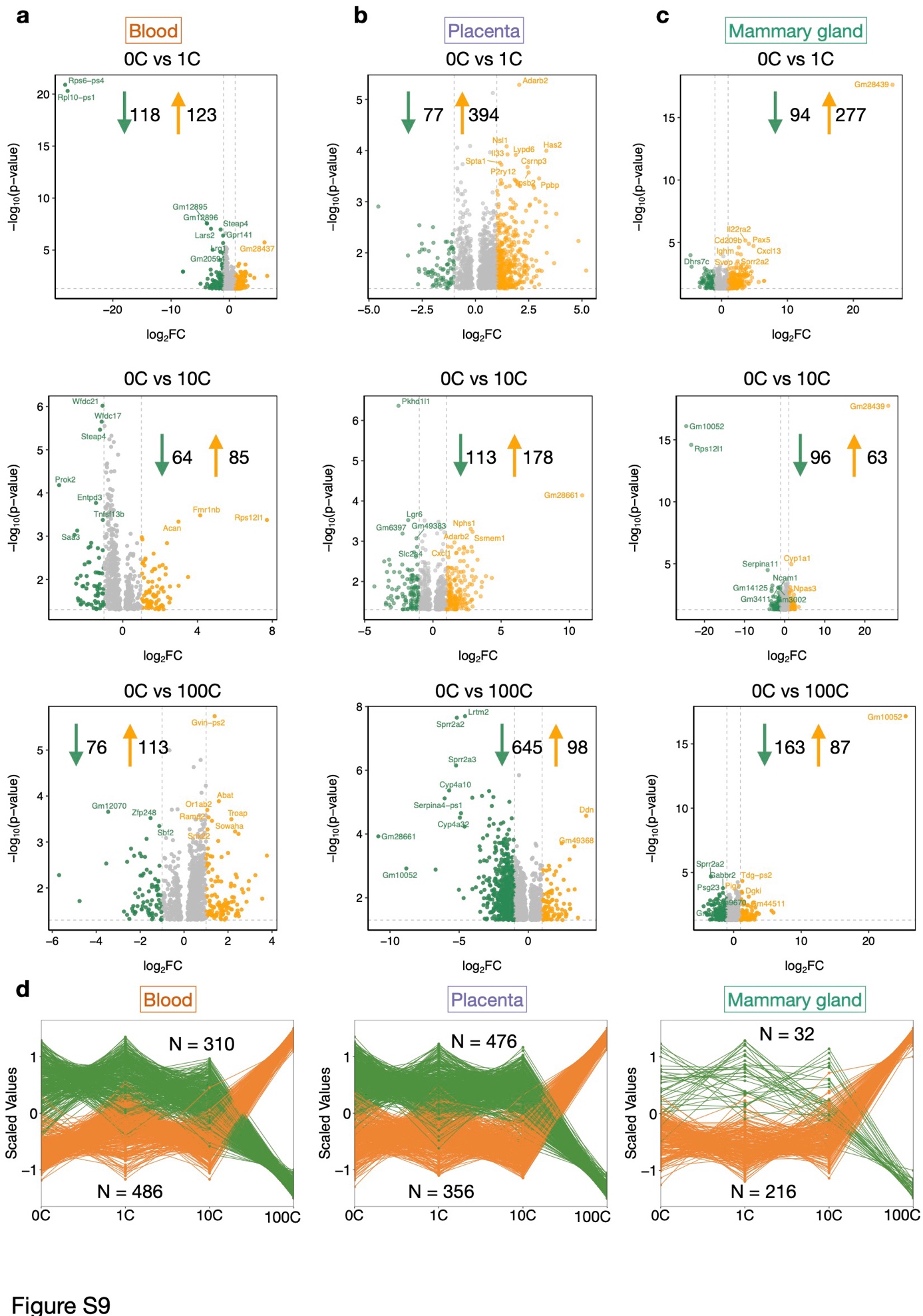

**Supplementary Fig. 9.** **Differentially expressed genes in response to MPs exposure.**

(a-c) Differential expression genes (DEGs) in blood (a), placenta (b), and mammary gland (c), across different dose groups compared to 0C.

(d) The increasing (orange) and decreasing (green) cluster of the genes in blood, placenta, and mammary gland, estimated by GLM (p < 0.01).

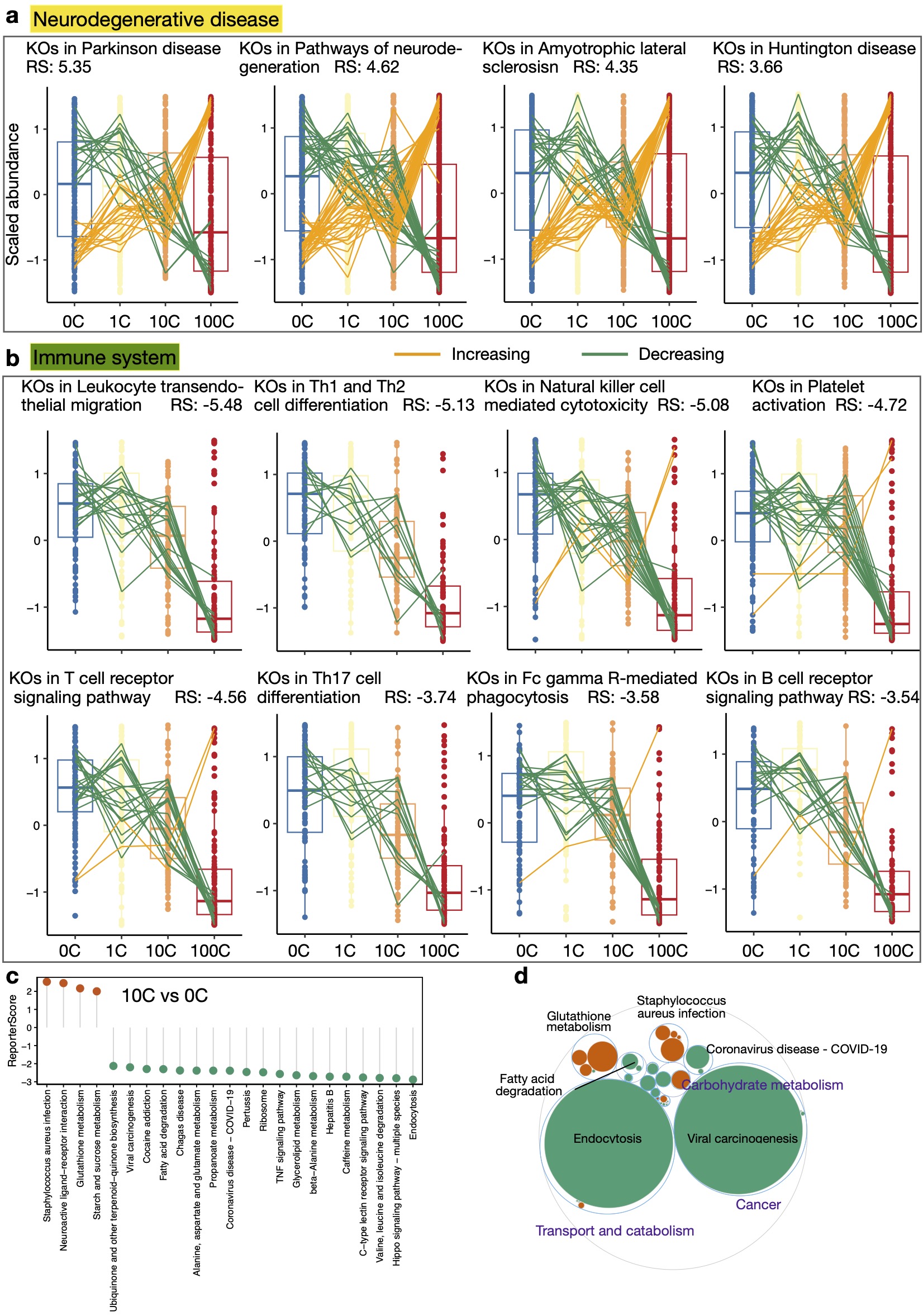

**Supplementary Fig. 10.** **Blood functional shifts in response to MPs exposure.**

(a-b) Box chart of KEGG pathways related to neurodegenerative disease (a) and immune system (b). Line colors indicate KO abundance trends. RS: ReporterScore.

(c) Lollipop chart of significantly enriched pathways (|ReporterScore| > 2.0, p < 0.05) in 0C and 10C groups.

(d) Gene-function integration maptree related to differential enriched pathways between 10C and 0C groups (|ReporterScore| > 2.0, p < 0.05), showcasing functional categories, KEGG pathways, and gene levels, with the size of filled circles representing average gene expression abundance in 10C and 0C groups and their differential trends (enriched in 10C in red, depleted in 10C in green).

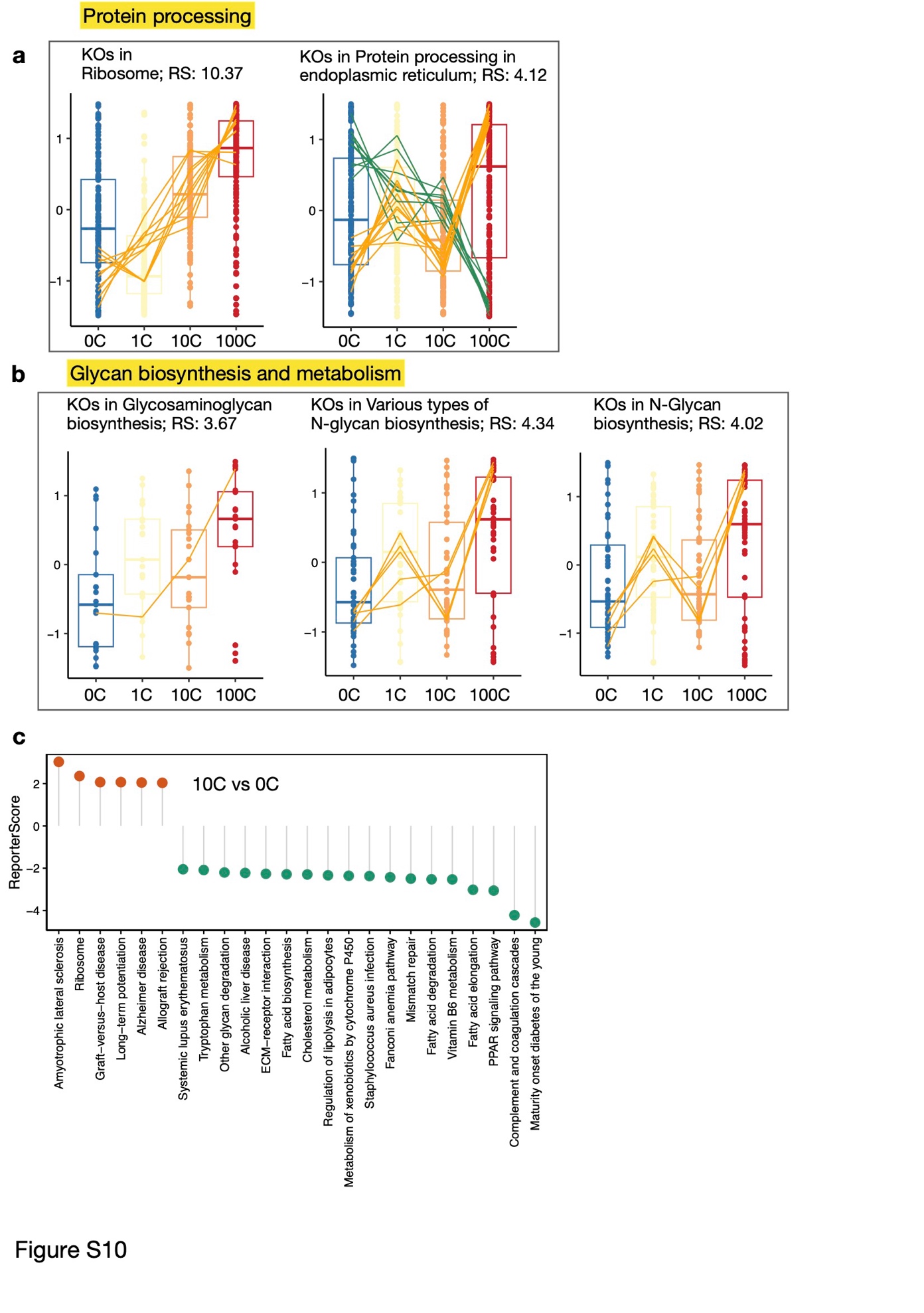

**Supplementary Fig. 11.** **Placental functional shifts in response to MPs exposure.**

(a-b) Box chart of KEGG pathways related to protein processing (a) and glycan biosynthesis and metabolism (b). Line colors represent KO abundance trends. RS: ReporterScore.

(c) Lollipop chart of significantly enriched pathways (|ReporterScore| > 2.0, p < 0.05) between 0C and 10C groups.

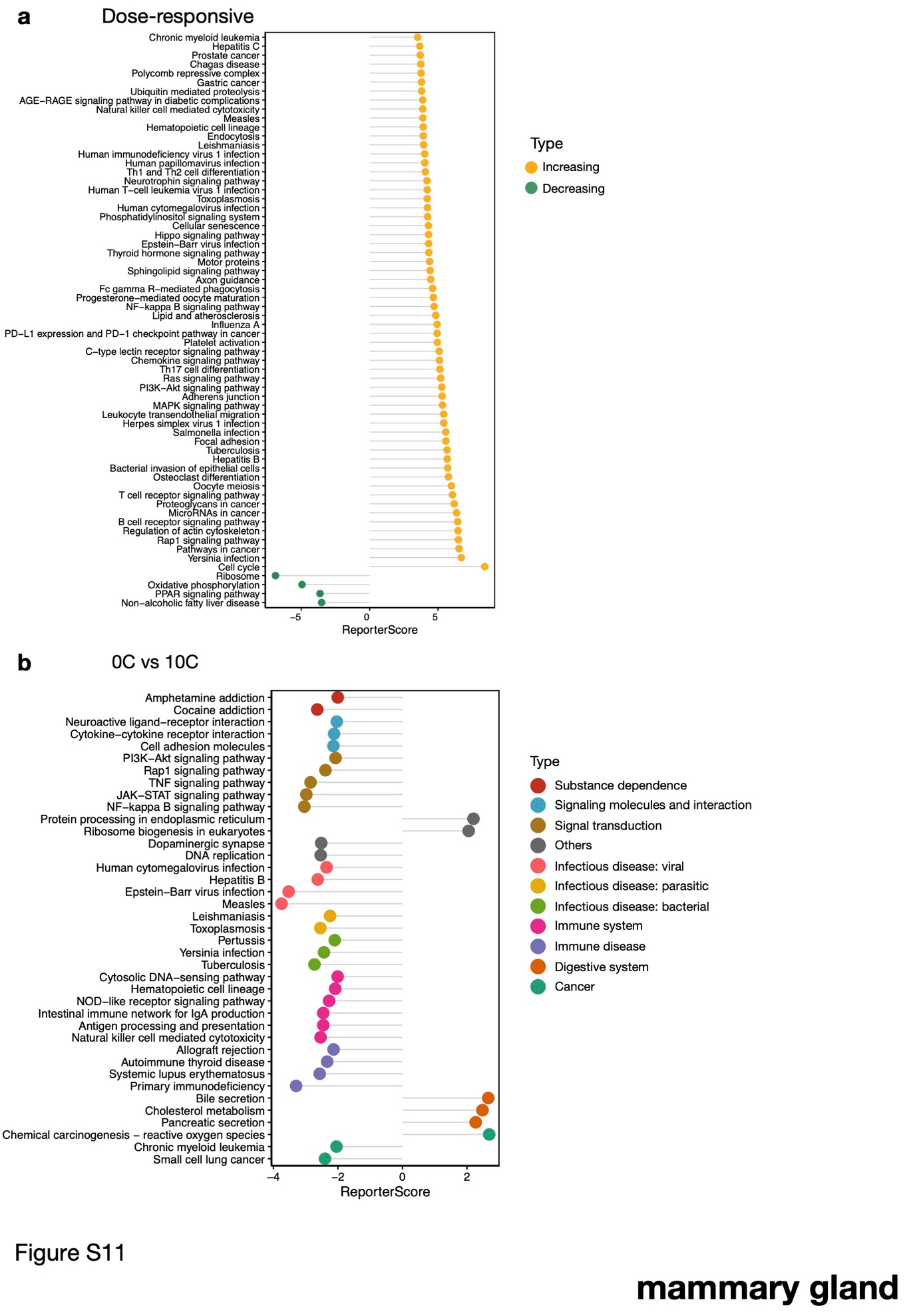

**Supplementary Fig. 12**. **Mammary gland functional dynamics in response to MPs exposure.**

(a) Lollipop charts displaying pathways with significant (|ReporterScore| > 3.5, p < 0.01) increases (yellow) or decreases (green) in reporter score.

(c) Lollipop chart of significantly enriched pathways (|ReporterScore| > 2.0, p < 0.05) between 0C and 10C groups.
